# CYP154C5 Regioselectivity in Steroid Hydroxylation Explored by Substrate and Protein Engineering

**DOI:** 10.1101/2020.03.04.972729

**Authors:** Paula Bracco, Hein J. Wijma, Bastian Nicolai, Jhon Alexander Rodriguez Buitrago, Thomas Klünemann, Agustina Vila, Patrick Schrepfer, Wulf Blankenfeldt, Dick B. Janssen, Anett Schallmey

## Abstract

CYP154C5 from *Nocardia farcinica* is a P450 monooxygenase able to hydroxylate a range of steroids with high regio- and stereoselectivity at the 16α-position. Using protein and substrate engineering based on the crystal structure of CYP154C5, an altered regioselectivity of the enzyme in steroid hydroxylation could be achieved. Thus, conversion of progesterone by mutant CYP154C5 F92A resulted in formation of the corresponding 21-hydroxylated product 11-deoxycorticosterone in addition to 16α-hydroxylation. Using MD simulation, this altered regioselectivity appeared to result from an alternate binding mode of the steroid in the active site of mutant F92A. MD simulation further suggested that water entrance to the active site caused higher uncoupling in this mutant. Moreover, exclusive 15α-hydroxylation was observed for wild-type CYP154C5 in the conversion of 5α-androstan-3-one, lacking an oxy-functional group at C17. Overall, our data give valuable insight into the structure-function relationship of this cytochrome P450 monooxygenase for steroid hydroxylation.

## Introduction

Cytochrome P450 monooxygenases (P450s or CYPs) are hemoproteins carrying a heme *b* molecule covalently linked to a cysteine side chain.^[1]^ From a biocatalytic perspective, they are remarkable enzymes as they are able to catalyze the selective hydroxylation of non-activated carbon atoms using molecular oxygen.^[2,3]^ For the activation of molecular oxygen during the catalytic cycle, they require electrons – coming from NAD(P)H – which are in most cases delivered to the monooxygenase by additional redox partners.^[4]^ One of the most important industrial uses of these cytochrome P450 monooxygenases is their application in steroid synthesis in the pharmaceutical industry due to their remarkable selectivity in steroid hydroxylation. Well-known examples include the 11β-hydroxylation of 11-deoxycortisol (Reichstein S) to hydrocortisone by *Curvularia* sp. or the conversion of progesterone to cortisone by *Rhizopus* sp.^[5–8]^ Though P450s are already applied on industrial scale, there is always the need for yield improvement, including the increase in hydroxylation specificity (*e.g*. fewer by-products), the need for an altered selectivity of the enzyme to hydroxylate e.g. also new sites in a known substrate or the adaptation of a known P450 to a new substrate.^[9]^ In that respect, protein engineering has proven to be a powerful tool to alter enzyme characteristics such as activity and selectivity. One example, reported by Kille *et al.*, is the generation of P450BM3 mutants hydroxylating testosterone selectively either at 2β- or 15β- position, while the starting mutant, P450BM3 F87A, forms a 1:1 mixture of 2β- and 15β- hydroxytestosterone.^[10]^ In this case, mutants were generated by iterative combinatorial active-site saturation mutagenesis, a protein engineering approach that is significantly facilitated by the availability of structural information for a given protein.

Recently, the crystal structure of CYP154C5, a cytochrome P450 monooxygenase from *Nocardia farcinica*, was elucidated in the presence of four steroid substrates.^[11]^ This enzyme catalyzes the highly regio- and stereoselective hydroxylation of different pregnans and androstans producing exclusively the corresponding 16α-hydroxylated products.^[12]^ As the natural redox partners of this P450 monooxygenase are unknown, putidaredoxin (Pdx) and putidaredoxin reductase (PdR) from *Pseudomonas putida* can be applied in bioconversions to supply CYP154C5 with electrons. The active site pocket of CYP154C5 forms a hydrophobic “tube” with two opposite polar regions at both ends. These polar regions are occupied by glutamine 239 and glutamine 398 forming hydrogen bond interactions with the hydroxyl or ketone functionalities of steroids at positions C3 (via water molecules) and C17.^[11]^ Additionally, several hydrophobic interactions between enzyme and steroid substrate within the active site could be identified. With the help of the crystal structure the remarkably high regio- and stereoselectivity of CYP154C5 towards steroids **1-6** (Figure 1) could also be explained.

**Figure 1.**
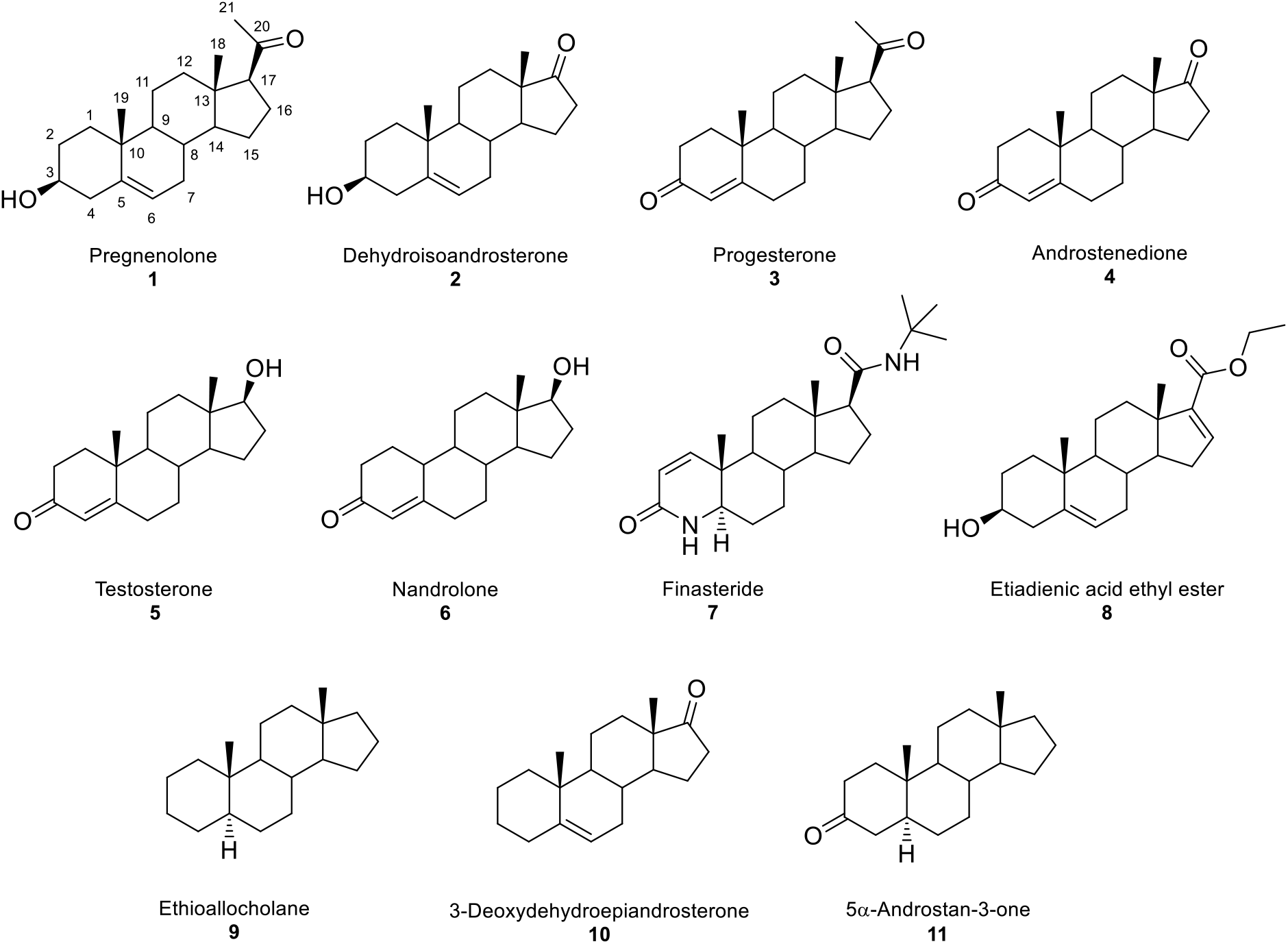
Chemical structures of previously tested (**1-6**)^[11,12]^ as well as five new steroid substrates (**7-11**) used in bioconversions with CYP154C5 (wild type and mutants), Pdx and PdR in this study.

With the future goal to modify the enzyme’s regioselectivity in steroid hydroxylation reactions, we herein aimed to explore the selectivity of CYP154C5 in more detail based on the previously obtained structural insight. To this end, selected active-site residues of CYP154C5, mediating important enzyme-substrate interactions, were mutated by site-directed mutagenesis, and resulting mutants were applied in bioconversions with steroids **1-6**. In a complementary approach, steroid substrates lacking specific functional groups, that otherwise enable hydrogen bonding with active site residues of CYP154C5, were tested in bioconversions with wild-type P450.

## Results and Discussion

### CYP154C5 mutagenesis

Based on the CYP154C5 crystal structure and a detailed analysis of the enzyme active site in the presence of different steroid substrates,^[11]^ four active-site residues have been identified to play an important role in steroid binding. Among them are the two glutamines at positions 239 and 398 forming hydrogen bonds with the carbonyl or hydroxyl groups at C3 (via a water molecule) and C17 of the steroid substrates (Figure 2). Moreover, residues M84 and F92 are involved in hydrophobic interactions with the steroid backbone.^[11]^ Thus, residues M84, F92, Q239 and Q398 were selected for mutagenesis to determine their role in steroid binding and CYP154C5’s selectivity in steroid conversions. To this end, single alanine mutations were prepared using site directed mutagenesis. Afterwards, wild-type CYP154C5 as well as mutants M84A, F92A, Q239A and Q398A were heterologously produced in *E. coli* C43 (DE3) and purified by anion exchange and affinity chromatography (Figure S1). Similarly, redox partners Pdx and PdR from *P. putida,* which are required for P450 activity, were produced in *E. coli* C43 (DE3) and subsequently purified by anion exchange and hydrophobic interaction chromatography (Figure S1).

**Figure 2.**
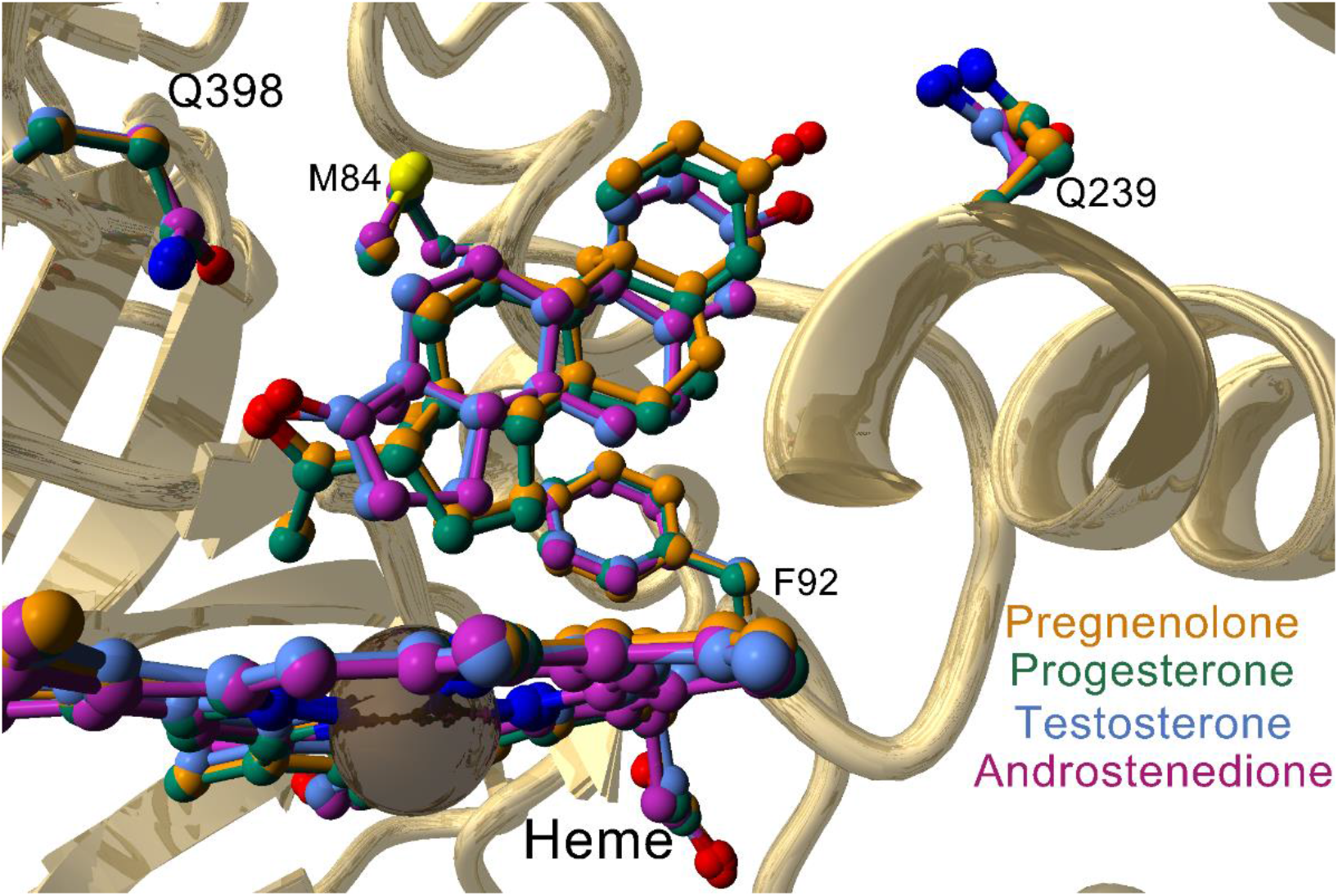
Superposition of the active sites of the four CYP154C5-steroid complexes (PDB codes: 4J6B, 4J6C, 4J6D and 4JBT) with the four amino acid residues M84, F92, Q239 and Q398 involved in steroid binding.

### Steroid conversion by CYP154C5 mutants

In order to analyze the influence of the different mutations on steroid binding and catalysis by CYP154C5, dissociation constants (K_D_), turnover numbers (TONs) and coupling efficiencies of the mutants towards steroids **1-6** were determined. In the absence of substrate, P450 enzymes exhibit an absorbance maximum around 420 nm. Upon substrate binding, a water molecule is removed as the sixth ligand at the heme iron.^[13]^ This causes a shift of the heme iron to a high-spin (HS) configuration which, in type I spectral changes, involves a shift of the P450 absorbance maximum to approximately 390 nm. From the peak-to-trough difference in absorbance (ΔA) between P450 with high spin iron (A_390_) and low spin iron (A_420_) dissociation constants can be inferred. Using the four different CYP154C5 mutants and steroids **1-6**, K_D_ values for almost all combinations could be determined (Table 1; respective plots are displayed in Figures S2-S6). Some mutants, however, yielded only a partial or no spectral shift upon substrate addition (Figure S7), even at high steroid concentration, hampering K_D_ determination.

**Table 1.** Dissociation constants (K_D_) of CYP154C5 wild type and mutants for steroid substrates **1-6**. K_D_ values were determined at 30°C for the His-tagged CYP154C5 mutants using the quadratic tight-binding equation.^[14]^ All values are the result of duplicate measurements.

| Substrate | $K_D$ (nM) | | | | |
| --- | --- | --- | --- | --- | --- |
|  | CYP154C5 | CYP154C5 M84A | CYP154C5 F92A | CYP154C5 Q239A | CYP154C5 Q398A |
| Pregnenolone (**1**) | 62 ± 14 | – [a] | –/+ [b] | <1 | 65 ± 21 |
| Dehydroepiandrosterone (**2**) | <1 | –/+ [b] | 1017 ± 32 | <1 | 687 ± 43 |
| Progesterone (**3**) | 15 ± 10 | –/+ [b] | 209 ± 12 | <1 | 34 ± 21 |
| Androstenedione (**4**) | 12 ± 7 | 3130 ± 234 | 512 ± 17 | <1 | 572 ± 151 |
| Testosterone (**5**) | 18 ± 10 | 5710 ± 161 | 1481 ± 33 | <1 | 747 ± 45 |
| Nandrolone (**6**) | 100 ± 17 | – [a] | –/+ [b] | 13 ± 6 | – [a] |

[a] No shift to high-spin heme observed.
[b] Only partial shift to high-spin heme observed.

Notably, obtained K_D_ values of the alanine mutants were in many cases higher compared to wild type CYP154C5, indicating a substantial influence of the mutated active-site residues on steroid binding. The main exception is CYP154C5 Q239A for which the obtained K_D_ values were even lower than for wild-type CYP154C5. This suggests a positive influence of mutation Q239A on steroid binding, possibly due to a loss of the hydrogen bonding via a water molecule with the oxygen atom at C3 of the steroids. Moreover, binding of pregnenolone (**1**) and progesterone (**3**) to CYP154C5 was hardly affected by amino acid exchange Q398A, whereas significantly higher K_D_ values were obtained for all other steroid substrates. Removal of the glutamine side chain in CYP154C5 Q398A leads to a loss of a hydrogen bond to the C17-substituent oxygen of the steroid substrate, and generates space that could result in a possible movement of steroids **2**, **4**, **5** and **6** in the active site. In contrast, steroids **1** and **3** carry a more spacious acetyl side chain at C17, which could fill this space resulting in a tighter binding. Interestingly, mutation M84A in CYP154C5 affected substrate binding the most among all tested variants. More than 100-fold lower binding affinities compared to wild-type CYP154C5 were obtained with androstenedione (**4**) and testosterone (**5**). Additionally, no spectral shifts of CYP154C5 M84A were observed with steroids **1** and **6**, while substrates **2** and **3** induced only partial shifts.

Additionally, also turnover numbers and coupling efficiencies of the purified mutants were determined in the conversion of steroids **1-6**, together with Pdx and PdR as redox partners (Tables 2 and 3). As a result, mutants CYP154C5 M84A, CYP154C5 F92A and CYP154C5 Q398A exhibited significantly decreased TONs in comparison to the wild-type enzyme, independent of the used substrate, while for mutation Q239A the TONs were less affected (Table 2). This is in general agreement with the observed higher K_D_ values obtained for mutants CYP154C5 M84A, CYP154C5 F92A and CYP154C5 Q398A. Interestingly, CYP154C5 M84A converted pregnenolone (**1**) to a small extent even though no shift from low to high spin could be detected. Such a behavior was previously observed in the case of cytochrome P450 from *Bacillus megaterium,* CYP106A2, where no change in absorbance was obtained upon addition of deoxycortisone (DOC) although this substrate is converted by this enzyme producing 15β-hydroxydeoxycortisone.^[15]^ Simgen *et al*., however, proved by FT infrared spectroscopy using the stretch vibration of the heme iron CO-ligand that DOC indeed enters the active site and binds close to the heme prosthetic group.^[15]^

**Table 2.** Turnover numbers (TON) of CYP154C5 mutants in the conversion of steroids **1-6**. Measurements were performed in duplicate.

| Substrate | TON (min<sup>-1</sup>) |  |  |  |  |
| --- | --- | --- | --- | --- | --- |
|  | CYP154C5<sup>[a]</sup> | CYP154C5 M84A | CYP154C5 F92A | CYP154C5 Q239A | CYP154C5 Q398A |
| Pregnenolone (**1**) | 4.47 ± 0.36 | 0.20 ± 0.00 | 0.85 ± 0.16 | 2.48 ± 0.02 | 0.93 ± 0.14 |
| Dehydroepiandrosterone (**2**) | 5.1 ± 0.11 | 0.75 ± 0.09 | 1.48 ± 0.26 | 2.07 ± 0.24 | 1.22 ± 0.21 |
| Progesterone (**3**) | 5.7 ± 0.22 | 2.52 ± 0.12 | 1.02 ± 0.07 | 4.33 ± 0.05 | 1.45 ± 0.07 |
| Androstenedione (**4**) | 3.14 ± 0.36 | 1.53 ± 0.14 | 1.68 ± 0.07 | 2.42 ± 0.05 | 1.77 ± 0.19 |
| Testosterone (**5**) | 1.77 ± 0.06 | 0.70 ± 0.05 | 1.75 ± 0.12 | 1.32 ± 0.12 | 0.57 ± 0.09 |
| Nandrolone (**6**) | 1.33 ± 0.10 | – <sup>[b]</sup> | 1.33 ± 0.09 | 1.28 ± 0.02 | – <sup>[b]</sup> |

<sup>[a]</sup> Data taken from [11] for comparison<sup>[b]</sup> No conversion observed**Table 3.** Coupling efficiencies of CYP154C5 mutants in the conversion of steroids **1-6**. Measurements were performed in duplicate.

**Table 3.** Coupling efficiencies of CYP154C5 mutants in the conversion of steroids **1-6**. Measurements were performed in duplicate.

| Substrate | Coupling efficiency (%) |  |  |  |  |
| --- | --- | --- | --- | --- | --- |
|  | CYP154C5<sup>[a]</sup> | CYP154C5 M84A | CYP154C5 F92A | CYP154C5 Q239A | CYP154C5 Q398A |
| Pregnenolone (**1**) | 68 ± 1 | 34 ± 6 | 40 ± 8 | 27 ± 5 | 53 ± 2 |
| Dehydroepiandrosterone (**2**) | 86 ± 5 | 60 ± 7 | 55 ± 0 | 90 ± 4 | 41 ± 6 |
| Progesterone (**3**) | 84 ± 3 | 34 ± 1 | 22 ± 9 | 38 ± 8 | 46 ± 0 |
| Androstenedione (**4**) | 83 ± 1 | 23 ± 14 | 64 ± 3 | 66 ± 6 | 51 ± 7 |
| Testosterone (**5**) | 62 ± 5 | 39 ± 4 | 46 ± 3 | 64 ± 8 | 68 ± 1 |
| Nandrolone (**6**) | 40 ± 7 | – <sup>[b]</sup> | 52 ± 18 | 34 ± 3 | – <sup>[b]</sup> |

<sup>[a]</sup> Data taken from [11] for comparison<sup>[b]</sup> No conversion observed

Similarly, obtained coupling efficiencies of the CYP154C5 mutants in the conversion of steroids **1-6** were, in many cases, also negatively affected (Table 3). This is especially evident for substrates pregnenolone (**1**) and progesterone (**3**) as well as for coupling efficiencies of mutant CYP154C5 M84A with all tested steroids. In contrast, coupling efficiencies of CYP154C5 F92A with nandrolone (**6**), CYP154C5 Q239A with **3**, **5** and **6** as well as coupling efficiencies of CYP154C5 Q398A with testosterone (**5**) are still similar to the wild-type values.

In almost all cases, the CYP154C5 mutants still formed the corresponding 16α-hydroxylated products in steroid conversions of **1-6**. Hence, regioselectivity of the single mutants was generally not affected, except for the conversion of **3** by mutant CYP154C5 F92A. Here, formation of a second hydroxylation product was observed (Figure S12), which was identified as 11-deoxycorticosterone (hydroxylation at position 21) by NMR analysis. Both products, 16α- and 21-hydroxylated **3**, were produced in a ratio of 4:1 by CYP154C5 F92A. Hydroxylation of **3** as well as 17α-hydroxyprogesterone at C21 yielding 11-deoxycorticosterone and 11-deoxycortisol, respectively, are important steps in adrenal steroidogenesis required for the synthesis of glucocorticoids and mineralocorticoids. In human, CYP21A2 is the responsible enzyme catalyzing this step. Interestingly, for CYP21A2 the formation of 16α-hydroxyprogesterone as side product has been reported as well.^[16]^ Moreover, exchange of valine 359 by alanine yielded a mutant with significantly increased hydroxylation in 16α position resulting in a ratio of 21-hydroxyprogesterone to 16α-hydroxyprogesterone of 60:40, while mutant CYP21A2 V359G gave even 90% 16α-hydroxyprogesterone.^[17]^

Usually, hydroxylation of a methyl group (primary carbon such as C21 in **3**) is energetically less favored than hydroxylation of a methanediyl group (secondary carbon such as C16 in **3**) due to a significant difference in the energy barrier of oxidation.^[18]^ To reveal the structural basis of the change in regioselectivity of mutant CYP154C5 F92A in progesterone hydroxylation, this enzyme-substrate complex was studied by computational tools.

### Modeling of CYP154C5 F92A

A structural model of the compound I intermediate of the F92A mutant was generated and used in docking and molecular dynamics simulations. As a reference, a crystal structure of wild-type CYP154C5 with progesterone (**3**) bound was converted to the compound I state (Figure 3-A) and subjected to docking as well.^[11]^ The docking simulations with the F92A mutant suggested two possible binding orientations for substrate **3**. One orientation is similar to the one found in the crystal structure (Figure 3-C), whereas the second is clearly different: progesterone is reoriented with its A and B ring now occupying the space created by the F92A mutation and only the hydrogens of carbon 21 are close to the reactive oxygen of compound I (Figure 3-B). This alternative orientation of the substrate would cause extreme steric hindrance, if F92 would remain in the position that is observed in all X-ray structures; the aromatic ring of F92 would have to interlock with the A or B ring of the steroid.

**Figure 3.**
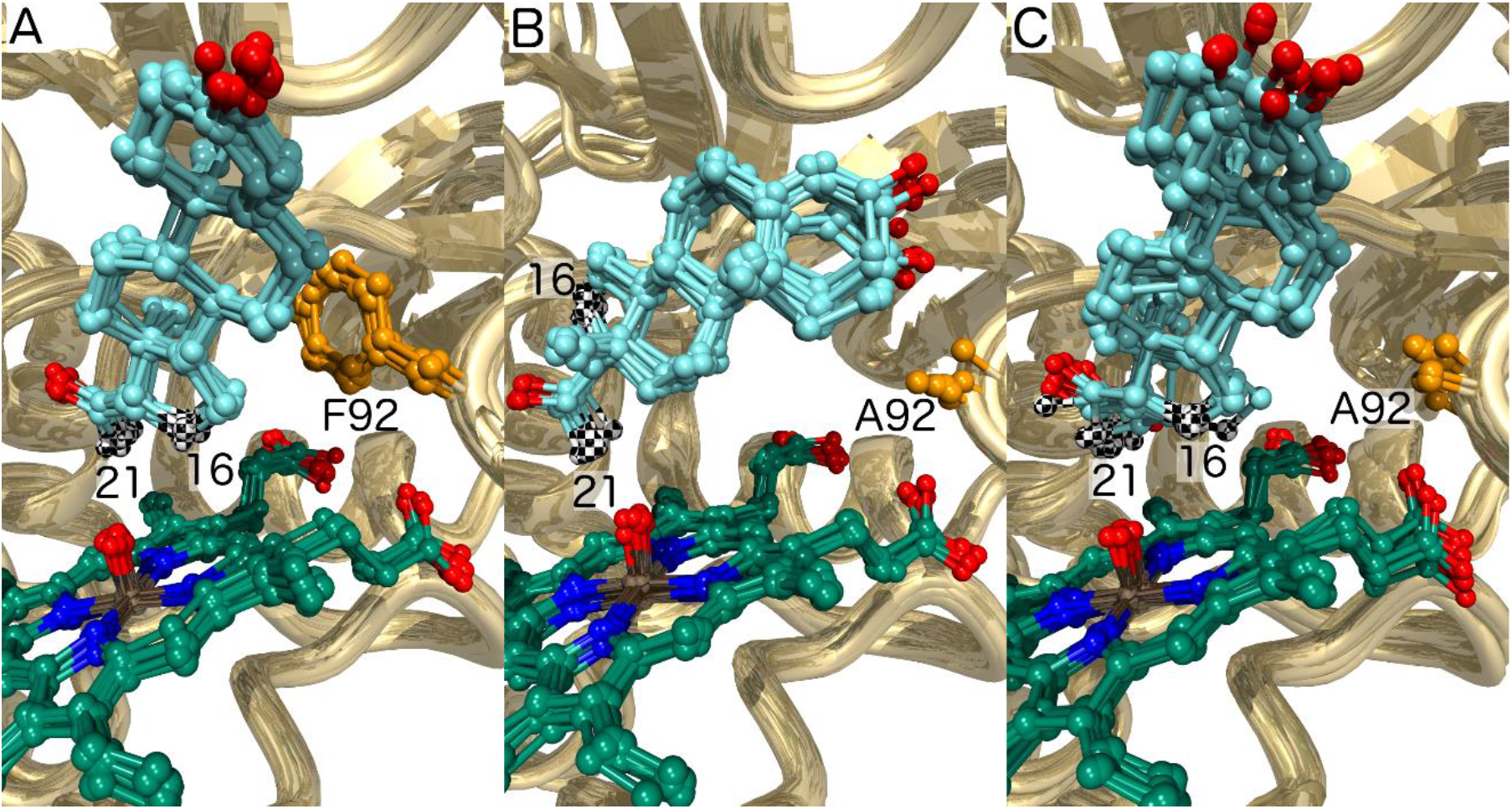
The modeled orientations of progesterone (**3**) can explain selective oxidations at the 21 and 16α positions. For each structure, 10 MD snapshots are shown. Carbon 16 and 21 of the progesterone (**3**) substrate are indicated with a black-white checker pattern. **A**) Wild type CYP154C5, where both the hydrogens at the 21 and 16α position are close to the compound I oxygen, suggesting the result will be selective oxidation of the more reactive secondary carbon atom 16 instead of the primary carbon atom 21; **B**) F92A with progesterone (**3**) bound in an alternative orientation, in which only oxidation at the 21 position is possible. **C**) F92A is also predicted to occur with progesterone (**3**) bound in a wild type-like orientation, enabling 16α oxidation.

The first step in P450-catalyzed hydroxylation is the abstraction of a hydrogen atom from the substrate by the electrophilic oxygen of the compound I intermediate.^[19]^ The predicted reactivity of different enzyme-substrate complexes was therefore examined by molecular dynamics (MD) simulations with scoring of near-attack conformations (NACs) in which substrate hydrogens approach the oxygen of the compound I intermediate (Figure S8). For each complex, three independent 22 ns MD simulations were performed. The enzyme-substrate complexes were stable (Figure 4) and the simulations gave reproducible results. In simulations of progesterone (**3**) in complex with wild-type CYP154C5, both the 16α and 21 hydrogen atoms stayed close to the reactive oxygen atom (Figure 4-A). The overall X-ray structure was maintained and high percentages of NACs were found for both positions (Table 4). The observation that only 16α hydroxylation took place with the wild-type enzyme is in agreement with the higher reactivity of this secondary carbon atom compared to the primary carbon atom at position 21,^[18]^ even though the latter might be influenced by the flanking carbonyl group.

**Table 4.** Near attack conformations of wild-type (WT) CYP154C5 and mutant F92A with progesterone as ligand. Potential oxidation sites of progesterone are given.

| Structure simulated by molecular dynamics | NACs during MD simulation (%) |  |
| --- | --- | --- |
| | 16α oxidation | 21 oxidation |
| WT + progesterone | 22 | 7 |
| F92A + progesterone in alternative orientation<sup>[a]</sup> | 0 | 42 |
| F92A + progesterone in native like orientation<sup>[b]</sup> | 2 | 5 |

<sup>[a]</sup> Also 0.4% NACs were observed for 12 $\beta$ oxidation.
<sup>[b]</sup> Additionally, 0.3% NACs for oxidation at position 17 was recorded.

**Figure 4.**
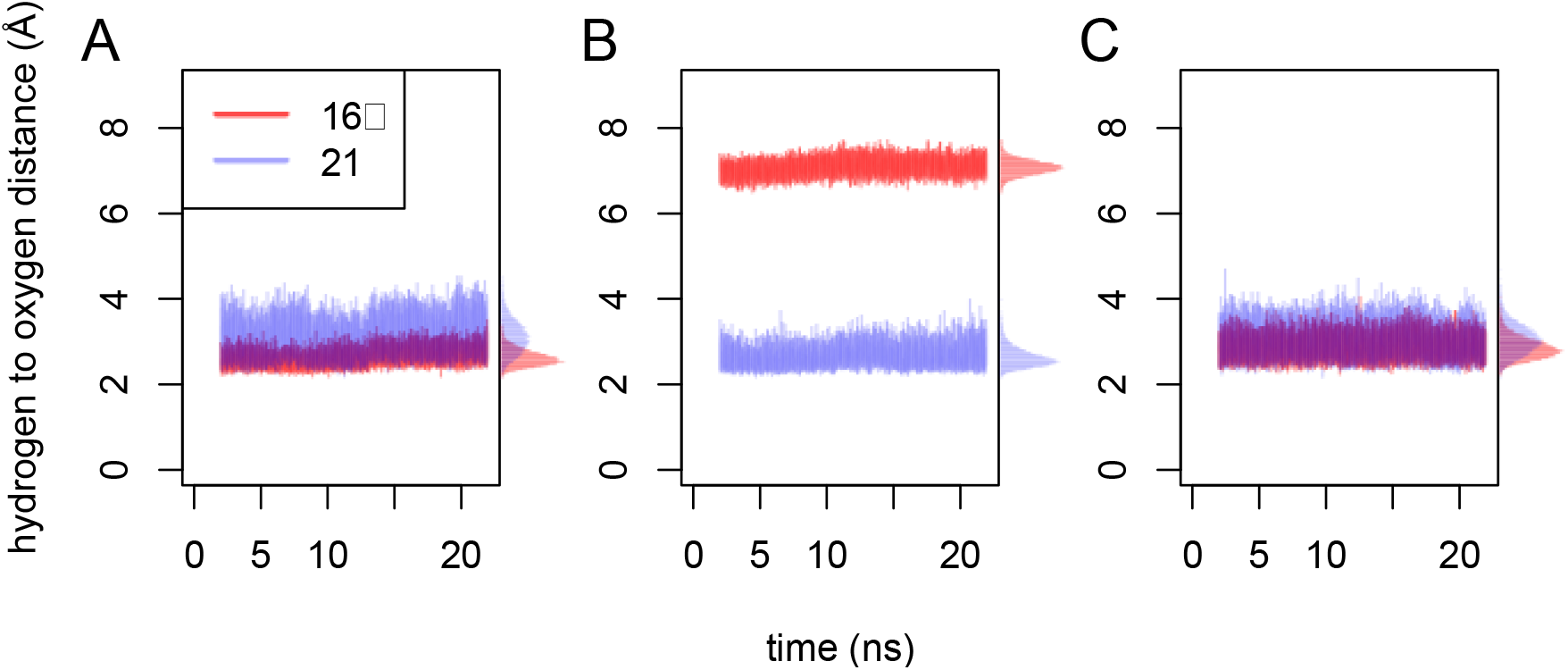
Distance of the substrate‘s hydrogen atoms to the oxygen of compound I. Because there are three identical 21 hydrogen atoms per substrate, only the shortest distance is shown at each time-point. Also for clarity, for each modeled enzyme substrate complex only the first out of the three independent MD simulations is shown. **A**) Wild-type CYP154C5 with progesterone (**3**) bound. **B**) CYP154C5 F92A with progesterone (**3**) bound in the alternative orientation; **C**) CYP154C5 F92A with progesterone (**3**) bound in the native-like orientation.

With the substrate oriented in the F92A mutant like it is in the wild-type enzyme, the distances and NAC percentages for the 16α and 21 hydrogens calculated from the simulations were similar to those found with the wild-type CYP154C5 (Figure 4-C). In contrast, MD simulations of the F92A mutant with progesterone (**3**) bound in the alternative orientation (Figure 4-B) showed that in this case only the 21 position can undergo oxidation. The distance between the 16α hydrogen and the reactive oxygen is predicted to exceed 6 Å, making hydrogen abstraction impossible and only the 21 hydrogens gave significant levels of NACs (Table 4). Thus, the modeling suggests that the F92A mutation provides an additional progesterone (**3**) binding mode that is particularly suitable for oxidation at the 21 position. Also Pallan *et al.* suggested that two alternate binding modes of **3** in CYP21A2’s active site are responsible for the formation of 21- and 16α-hydroxylated products by this enzyme based on an observed partial burst in pre-steady state kinetics.^[20]^

Furthermore, CYP154C5 F92A with **3** also gave the highest uncoupling (Table 3). Uncoupling is a common challenge of many P450 mutants which is highly relevant as it hinders their use in applied catalysis. In P450s, it can be explained by a poor enzyme-substrate structural complementarity, which allows water to interfere and react with the activated oxygen before the latter can abstract a hydrogen from the substrate.^[21]^ In agreement with this hypothesis, we found that substrate **3** is more mobile in the active site of mutant F92A. The root mean square fluctuations (RMSF) of this substrate are higher in the mutant than in wild type CYP154C5 (Figure S9). Also, while in case of the wild type the closest water stayed at a 7 Å distance from the reactive oxygen during the entire MD simulations, water molecules did approach the reactive oxygen for both substrate orientations in simulations with the F92A enzyme (Figure 5). Some of the observed distances are less than 2 Å; the intruding water makes Van der Waals contact with the reactive oxygen. Thus, the modeling is in agreement with the idea that higher substrate mobility causes uncoupling by allowing for water encroachment.

**Figure 5.**
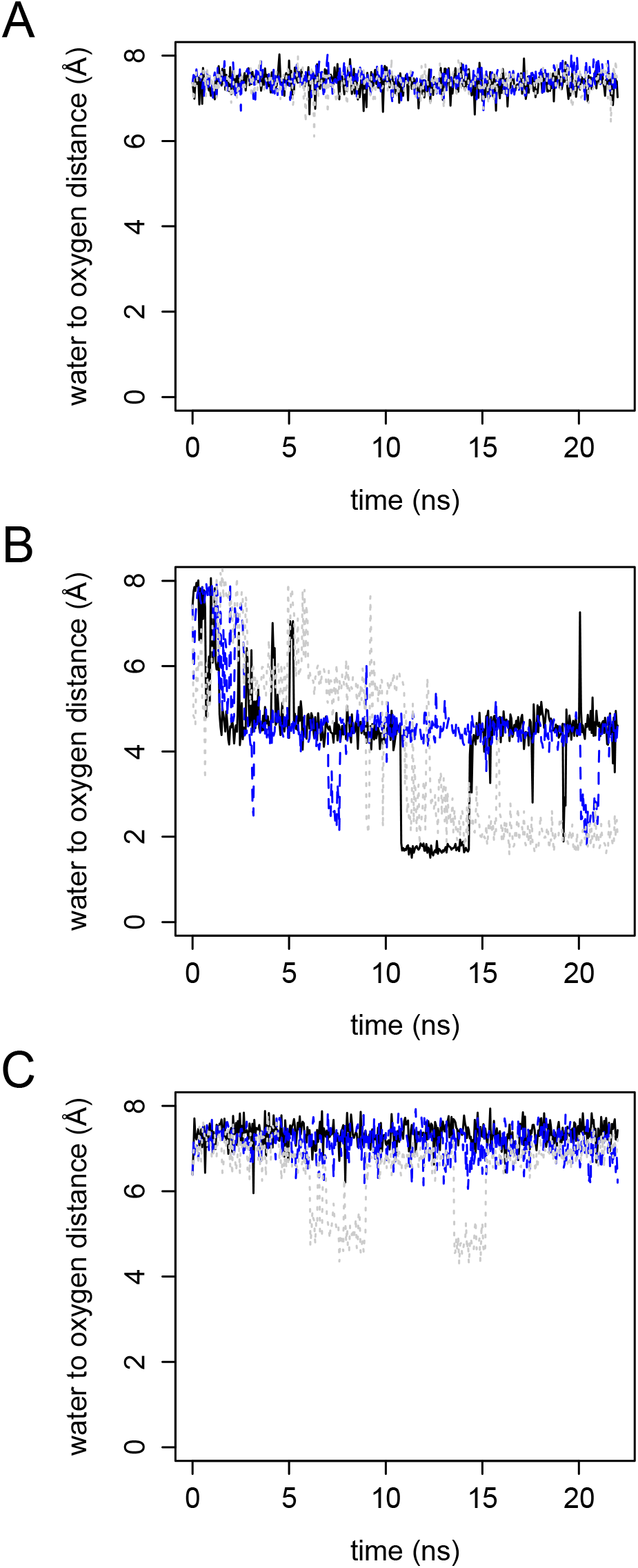
Distance between the reactive oxygen atom of compound I and the nearest water molecule during MD simulation. The results from three independent MD simulations are shown with a black continuous line (first simulation), with blue long dashes (2^nd^ simulation), and with gray short dashes (3^rd^ simulation). **A**) Wild type CYP154C5 with progesterone (**3**) bound. **B**) CYP154C5 F92A with substrate **3** bound in the alternative orientation; **C**) CYP154C5 F92A with substrate **3** bound in the native like orientation.

### Bioconversions of new steroid substrates by wild-type CYP154C5

Instead of introducing mutations, an alternative approach to study the enzyme’s selectivity is the selection of steroid substrates that are either lacking key functional groups for the enzyme-substrate interaction or that are carrying new features. Hence, based on the known CYP154C5-steroid complex structures, five new steroid substrates were selected. As previously reported, oxyfunctional groups at C3 and C17 were shown to form hydrogen bond interactions with residues Q239 (via a water molecule) and Q398, respectively.^[11]^ Therefore, steroid substrates lacking one (**10** and **11**) or both (**9**) oxyfunctional groups, as well as steroids containing a larger side chain at position C17 (**7** and **8**) were chosen to be tested in bioconversions with wild-type CYP154C5 (Figure 1).

Initially, small-scale reactions employing whole cells or cell-free extract of *E. coli* containing wild-type CYP154C5, Pdx and PdR were carried out in order to investigate if compounds **7-11** are converted. Reactions employing whole cells or cell free extract of *E. coli* containing only Pdx and PdR were used as negative controls under the same reaction conditions. As a result, no conversion of steroid substrates **7** and **8** by CYP154C5 could be observed, neither employing whole cells nor cell-free extract. This is probably the result of the larger side-chain at C17 preventing binding of the steroids in the enzyme’s active site. This is further supported by the fact that **7** and **8** did not induce any spectral shift during K_D_ measurements. Similarly, CYP154C3 from *Streptomyces griseus*, a homologue of CYP154C5 hydroxylating steroids selectively at 16α position as well, is also unable to convert steroids with bulky substituents at the D ring.^[22]^ In case of ethioallocholane (**9**) conversion by CYP154C5, a possible product peak was identified by GC-MS (Figure S13) though conversion was too low for product isolation. A NIST-library search suggested a steroid-related structure for this product. Additionally, steroid **9** induced a partial spectral shift of CYP154C5 during K_D_ measurements with a high-spin species content of roughly 50% (Figure S7-G). These results suggest that substrate **9** is indeed converted by CYP154C5 but further tests will be necessary to identify the formed product. In contrast, conversion of substrate **10** led to the formation of several products (Figure S14), indicating that the regio- and/or stereoselectivity of CYP154C5 was altered. Moreover, conversion of substrate **11** by CYP154C5 resulted in one hydroxylated product (Figure S18).

In order to elucidate the structures of formed products, whole-cell bioconversions were performed on preparative scale for substrates 3-deoxydehydroepiandrostendione (**10**) and 5α-androstan-3-one (**11**). Similar to the results obtained on analytical scale, several products were formed in the preparative-scale conversion of **10** by wild-type CYP154C5. Of these products, two could be obtained in sufficient amount and purity for subsequent NMR analysis (see supplementary information for respective NMR spectra). Results revealed that 16α-hydroxy-3-deoxydehydroepiandrostendione was formed as the main product. Interestingly, the second purified product seems to be the result of a double hydroxylation as indicated by GC-MS and NMR data (Figures S16 and S17). The exact hydroxylation positions, however, could not be determined due to low product quantities. Preliminary docking studies with **10** and the compound I model of wild-type CYP154C5 suggest position 4β as potential hydroxylation site in addition to the observed 16α-hydroxylation (Figure S10-A). Similarly, also product 16α-hydroxy-3-deoxydehydroepiandrosterone could be docked in the active site with position 4β as potential hydroxylation site (Figure S10-B), which would ultimately result in a double hydroxylation of **10**. Furthermore, position 2α was identified as potential additional hydroxylation site when using 4β-hydroxylated **10** as docking substrate (Figure S10-C). Even though these docking poses were only obtained *in silico*, they can give a first indication for potential hydroxylation sites of the other observed products.

In contrast, the product formed in the preparative-scale conversion of **11** by CYP154C5 was identified as 15α-hydroxy-5α-androstan-3-one by NMR (Figure S18). This result indicates a change in CYP154C5’s regioselectivity in the conversion of **11**, likely caused by the lack of the functional group at position C17 and/or the saturated A-ring of the steroid substrate. To gain structural insight into the altered regioselectivity of wild-type CYP154C5 in the conversion of **11**, the P450 was co-crystallized with this steroid. The overall structure of CYP154C5 is very similar to the previously determined CYP154C5 structures with Cα-RMSD values of 0.20-0.37. The electron density of the ligand and the derived model indicate that **11** binds in a similar position as steroids **4** and **5** (Figure 6), the latter being hydroxylated in 16α position by wild-type CYP154C5. Interestingly, however, C15 of steroids **4**, **5** and **11** in the corresponding crystal structures of CYP154C5 are in a similar position as C16 of steroids **1** and **3** (Figure 6 A), which are also hydroxylated in 16α position by wild-type CYP154C5. Hence, the crystal structure alone cannot explain the observed hydroxylation of **11** in 15α-position. Therefore, MD simulations (10 trajectories of each 22 ns) of the compound I state of the CYP154C5 structure with **11** bound in the active site have been performed with scoring of NACs as mentioned before (see Figure S11). As a result, predicted NAC percentages of 0.31% for C15 and 15.8% for C16 were obtained. This confirms that C15 can be reached in productive conformations by the oxygen of compound I, even though the probability for attack of C16 seems to be higher as judged by distance. The question, why only 15α-hydroxy-5α-androstan-3-one is observed as product in conversions of **11** by CYP154C5, cannot finally be solved but might be caused by a higher reactivity of C15 compared to C16. Here, quantum mechanical calculations could be performed in the future to reveal further insight.

**Figure 6.**
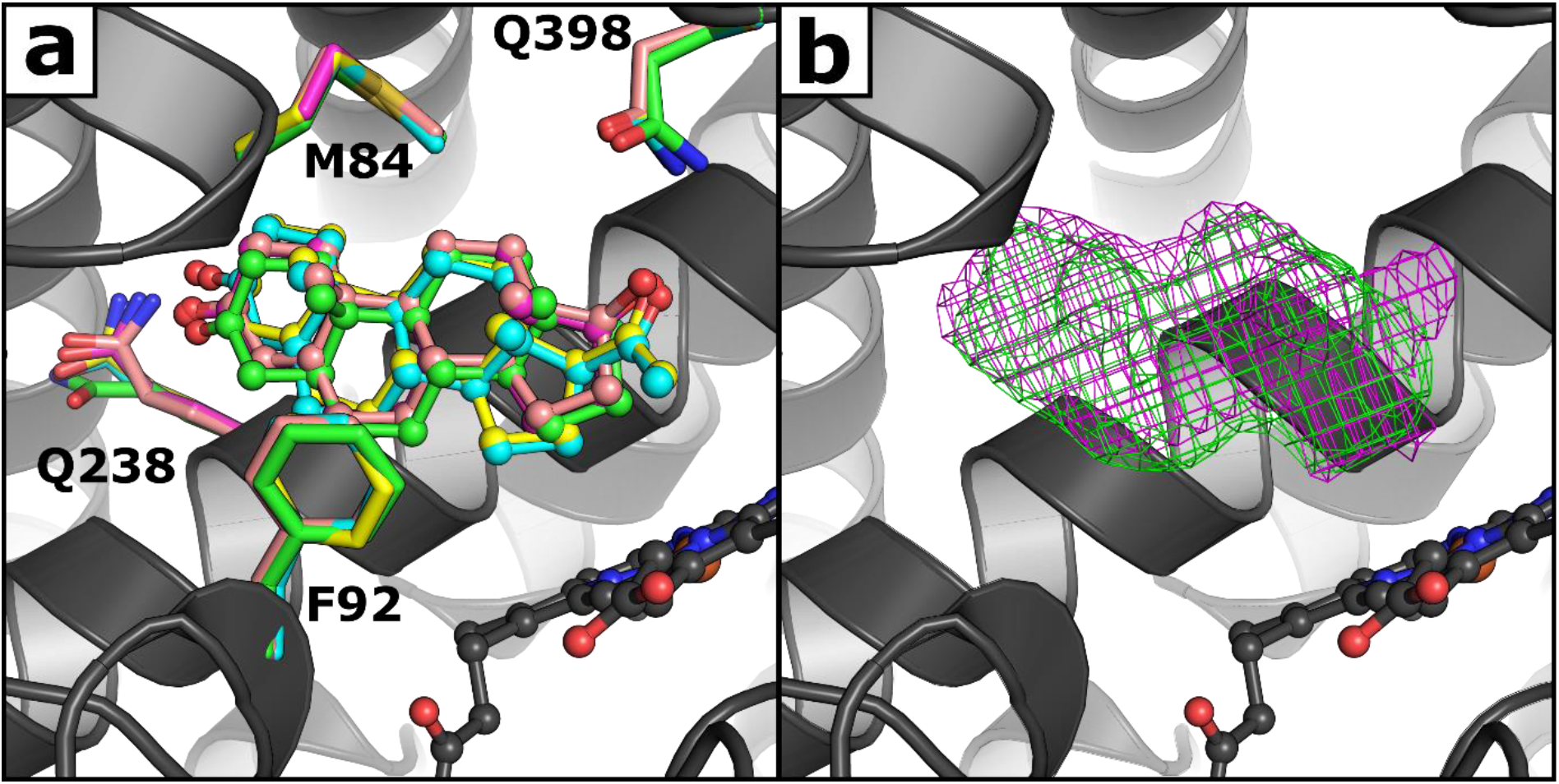
(a) Superimposition of the crystal structures of CYP154C with bound compound **11** (green) and the published steroid complexes [PDB-codes: 4J6B (**1**, yellow), 4J6C (**3**, cyan), 4J6D (**5**, pink) and 4JBT (**4**, magenta)]. (b) Mesh representing a 2F_o_F_c_ electron density contoured at a σ level of one carved around the steroids which are colored according to (a).

Additionally, K_D_ values, TONs and coupling efficiencies for wild-type CYP154C5 in the conversion of substrates **10** and **11** were determined. As a result, CYP154C5 exhibited a rather high affinity towards substrates **10** and **11** with K_D_ values of 94 ± 52 and 20 ± 16 nM, respectively. In contrast, coupling efficiencies are dramatically decreased in both cases resulting in only 7 ± 6 and 26 ± 10% for steroids **10** and **11,** respectively. Similarly, obtained TONs (0.67 ± 0.17 and 0.77 ± 0.17 min^−1^ for **10** and **11**, respectively) are also rather low compared to steroids **1-6**. This suggests that steroid conversion by CYP154C5 is indeed significantly affected if one of the oxyfunctional groups at C3 or C17 of the steroid backbone, and hence the corresponding hydrogen bond, is missing.

## Conclusion

With our study, we could demonstrate experimentally that the previously reported high regioselectivity of CYP154C5 is dependent on the presence of oxyfunctional groups at C3 and C17 of the steroid substrate, as products with new hydroxylation sites could be obtained in reactions with steroids lacking one of those substituents. Here, hydroxylation of **11** in 15α position is especially interesting, as 15α hydroxylation of steroids by bacterial cytochrome P450 monooxygenases has hardly been observed so far. In contrast, the replacement by alanine of active site residues Q239 and Q398, which have been shown to interact with the oxyfunctional groups through hydrogen bonding, did not alter the enzyme’s regioselectivity but had a negative impact on catalytic efficiency. Similarly, mutagenesis of residues M84 and F92, which form hydrophobic interactions with the steroid backbone, resulted in reduced turnover numbers and coupling efficiency as well as higher dissociation constants for most steroid substrates tested. This was especially evident for mutant M84A, which confirms the importance of residue M84 for delimiting the active site pocket and confining the steroid in a catalytically active position. Moreover, mutation F92A appeared to enable a second binding orientation of progesterone (**3**) in the enzyme active site resulting in C21 hydroxylation. Overall, our data demonstrate the feasibility for future modification of CYP154C5 regioselectivity by protein engineering and give valuable insight into the structure-function relationship of this cytochrome P450 monooxygenase for steroid hydroxylation.

## Experimental Section

### Substrates and chemicals

Pregnenolone (**1**), progesterone (**3**), testosterone (**5**) and cholesterol were purchased from Sigma-Aldrich (Taufkirchen, Germany). Dehydroepiandrosterone (**2**), androstenedione (**4**), nandrolone (**6**), finasteride (**7**), etiadienic acid ethyl ester (**8**), ethioallocholane (**9**), 3-deoxydehydroepiandrostendione (**10**) and 5α-androstan-3-one (**11**) were purchased from Steraloids Inc. (Newport, USA). Cytochrome c from horse heart, formate dehydrogenase from *Candida boidinii*, superoxide dismutase, catalase from bovine liver and hydroxypropyl β-cyclodextrine were purchased from Sigma Aldrich. All solvents and other chemicals required for the experiments were obtained from Roth (Karlsruhe, Germany) or Sigma-Aldrich and used without further purification.

### Bacterial strains and plasmids

*Escherichia coli* DH5α (Invitrogen, Carlsbad, CA, USA) was used for genetic manipulations, while *E. coli* C43 (DE3) (Lucigen, Middleton, WI, USA) was used for recombinant gene expressions. Plasmid pET28a(+) was purchased from Novagen (EMD Biosciences, San Diego, CA, USA). Preparation of plasmid pACYCcamAB for coexpression of putidaredoxin reductase (CamA or PdR) and putidaredoxin (CamB or Pdx) from *Pseudomonas putida* was described elsewhere.^[23]^ Preparation of plasmid pIT2cyp154c5 was described elsewhere.^[12]^ The gene of CYP154C5 was subcloned from vector pIT2cyp154c5 into pET28a(+) using restriction sites NdeI and HindIII. The resulting plasmid was named pET28cyp154C5. Expression of CYP154C5 from vector pET28cyp154C5 results in a fusion protein with N-terminal His-tag.

### Generation of CYP154C5 mutants

Mutants were prepared by QuikChange^®^ site-directed mutagenesis using the Pfu-Turbo Hotstart PCR Master Mix (Agilent, Waldbronn, Germany) according to the manufacturer’s instructions. The primers applied in the PCR reactions are listed in Table S1 in the supplementary. Correct introduction of the mutations was confirmed by sequencing at GATC Biotech (Konstanz, Germany) and final plasmids were transformed into *E. coli* C43 (DE3) for protein expression.

### Expression and purification of enzymes

Production of CYP154C5 wild type and its mutants using *E. coli* C43 (DE3), as well as expression of *camA* (PdR gene) and *camB* (Pdx gene) using *E. coli* C43 (DE3) (pACYCcamA) and *E. coli* C43 (DE3) (pACYCcamB), respectively, were performed as described elsewhere.^[11]^ Protocols for the production of *E. coli* C43 (DE3) (pIT2cyp154C5) (pACYCcamAB) and *E. coli* C43 (DE3) (pETcyp154C5-F92A) (pACYCcamAB) whole-cell biocatalysts were also previously described.^[12]^ Purification of wild-type CYP154C5 and its mutants via N-terminal His-tag, as well as purification of PdR and Pdx by anion exchange and hydrophobic interaction chromatography were also performed as previously described.^[11]^

### Enzyme assays

The P450 concentration was measured using CO-difference spectra.^[24]^ The activity of the purified electron transfer components (ETC) Pdx and PdR was determined by cytochrome *c* reduction assay, monitoring the increase in absorbance at 550 nm of reduced cytochrome c (*ε*_450_ = 19.1 mM^−1^ cm^−1^) in a mixture containing a ratio of PdR:Pdx equal to 3:16.^[25]^ Additionally, all enzymes assays were also measured with cell lysate of the respective *E. coli* C43 (DE3) cells containing P450, PdR and Pdx before whole-cell catalysis as described elsewhere.^[11]^ Total protein concentration was determined by Bradford assay.^[26]^

### Substrate binding studies

Dissociation constants (K_D_) of CYP154C5 wild type and CYP154C5 mutants for the different steroids were determined by spectroscopic measurements upon titration of purified P450 with increasing steroid concentrations.^[13]^ Thus, purified CYP154C5 carrying an N-terminal His-tag was diluted with 50 mM potassium phosphate buffer pH 7.4 in order to reach 3 µM final enzyme concentration. To this mixture, substrate was added in concentrations from 0 to 150 µM. Therefore, three different substrate stock solutions of 0.1, 0.5 and 1 mM in 0.1-4.5% (w/v) hydroxypropyl-β-cyclodextrin in deionized water (diH_2_O) were prepared. The absorbance spectra of each sample were measured on a Cary 50 spectrophotometer (Agilent, Waldbronn, Germany) between 300 and 500 nm at 30 °C. As a blank, 3 µM P450 in 50 mM potassium phosphate buffer, pH 7.4, with addition of an equivalent amount of buffer instead of substrate solution was used. Each sample was prepared and measured in duplicate. By plotting the resulting absorbance difference (Abs_386nm_ – Abs_420nm_) against the applied substrate concentration and fitting the data with the tight binding equation using MATLAB, K_D_ values for the different steroids were obtained.^[27,14]^

### Turnover number determination

Reactions for the determination of turnover numbers were carried out in 5 mL scale. Each reaction contained 3 µM P450, 3 µM PdR, 16 µM Pdx, 0.5 U mL^−1^ formate dehydrogenase from *Candida boidinii*, 150 mM sodium formate, 300 U mL^−1^ catalase from bovine liver, 50 µM NADH and 2 mM of the respective steroid substrate (**1-6**) in 50 mM potassium phosphate buffer pH 7.4. Steroid stock solutions of 4 or 5 mM concentration were prepared in 1.8-4.5% (w/v) hydroxypropyl-β-cyclodextrin in diH_2_O depending on the substrate. All bioconversions were carried out at 30 °C and 250 rpm for 20 h. During bioconversions, samples were taken at different time points for subsequent HPLC and GC analysis. For that, 0.25 mL of each reaction was extracted as described elsewhere.^[12]^ Turnover numbers were calculated based on substrate consumption and for the period of time where the highest substrate consumption rate was observed. Each reaction was performed in duplicate.

In contrast, in the case of steroids **10** and **11** TONs were determined based on whole-cell conversion. When performing preparative scale reactions of **10** and **11** using frozen cells of *E. coli* C43 (DE3) (pIT2cyp154C5) (pACYCcamAB), samples were taken over time and TONs were calculated as described in the previous paragraph.

### Coupling efficiency determination

NADH depletion during bioconversions of steroids by CYP154C5 mutants, Pdx and PdR was monitored in a spectrophotometer at 340 nm (*ε*_340_ =6.22 mM^−1^ cm^−1^). Reactions of 0.7 mL total volume included 0.4 µM purified P450, 0.4 µM purified PdR, 14 µM purified Pdx, 600 U mL^−1^ catalase from bovine liver, 200 U mL^−1^ superoxide dismutase, 200 µM NADH and 1 mM of the respective steroid (4 or 5 mM stock in 1.8-4.5% (w/v) of hydroxypropyl-β-cyclodextrin in diH_2_O) in 50 mM potassium phosphate buffer pH 7.4. After the NADH was completely consumed, 0.5 mL reaction mixture was extracted as described for the whole-cell catalysis and further analyzed by HPLC and GC in order to determine the conversion.

### Analytical-scale bioconversions

In case of whole-cell bioconversions, frozen cells of *E. coli* C43 (DE3) (pIT2cyp154C5) (pACYCcamAB) overexpressing Pdx, PdR and CYP154C5 were resuspended in 50 mM potassium phosphate buffer, pH 7.4, to the desired final OD_600_ of 40. All bioconversions were carried out in 1 mL scale at 30 °C and 250 rpm with addition of glucose (0.54 mg mL^−1^ final conc.) for cofactor regeneration and 1 mM substrate. Substrate stock solutions of steroids were prepared in 36% (w/v) hydroxypropyl-β-cyclodextrin in potassium phosphate buffer, pH 7.4. In detail, stocks with a final concentration of 2.5; 3.2; 4.2; 4.1 and 4.2 mM for substrates finasteride (**7**), etiadienic acid ethyl ester (**8**); ethioallocholane (**9**); 3-deoxydehydroepiandrostendione (**10**) and 5α-androstan-3-one (**11**) were prepared, respectively. Control reactions were carried out in parallel with *E. coli* C43 (DE3) (pACYCcamAB) containing only Pdx and PdR. After 20 hours of reaction, the bioconversions were extracted for subsequent HPLC and GC analysis. For that, 500 µL of sample was extracted twice with ethyl acetate (300 µL) and once with chloroform (250 µL). The organic phases were combined, dried with sodium sulfate and the solvent was removed under reduced pressure. As an exception, conversions performed with steroid **8** were acidified with 2 M HCl previous to the extraction procedure.

### Preparative-scale bioconversions

For the conversion of **10** and **11**, preparative-scale bioconversions were carried out in 50 mM potassium phosphate buffer, pH 7.4, using resting whole cells of *E. coli* C43 (DE3) (pIT2cyp154c5) (pACYCcamAB) at 30 °C and 250 rpm with the addition of glucose (0.54 mg mL^−1^) for cofactor regeneration. In case of substrate **10**, resting cells were resuspended in 100 mL buffer to OD_600_ ~ 40, equivalent to 7.8 µM CYP154C5 and an ETC activity of 8.1 U mL^−1^ (1.1 U mg^−1^ of total protein), as determined by CO-difference spectra and cytochrome c assay, respectively. Similarly, conversion of substrate **11** was performed using resting cells resuspended in 100 mL buffer to OD_600_ ~ 60, equivalent to 4.2 µM CYP154C5 and an ETC activity of 13.8 U mL^−1^ (1.4 U mg^−1^ of total protein). Initial substrate concentrations of 1 mM (**10** and **11**) were used; for that, stock solutions of substrates **10** and **11** were prepared in 36% (w/v) hydroxypropyl-β-cyclodextrin (in diH_2_O) and DMSO, respectively. Preparative-scale bioconversions of progesterone (**3**) were carried out in 50 mM potassium phosphate buffer, pH 7.4, in shake flasks using resting whole cells of *E. coli* C43 (DE3) (pIT2cyp154c5_F92A) (pACYCcamAB) at 30 °C and 250 rpm. In this case, cells were resuspended in 80 mL buffer to OD_600_ ~ 40 equivalent to 8 µM CYP154C5 and an ETC activity of 4.7 U mL^−1^ (0.6 U mg^−1^ of total protein), as determined by CO-difference spectra and cytochrome c assay, respectively. After 24 h of reaction, the complete reaction volume was extracted twice with ethyl acetate (50 and 40 mL) and once with chloroform, (30 mL), the organic phases were combined, dried with sodium sulfate and the solvent was removed under reduced pressure. Hydroxylated steroid products were afterwards purified by silica gel column chromatography with a mixture of ethyl acetate: *n*-heptane (8:2) as mobile phase.

### GC and HPLC analyses

In the case of pregnenolone (**1**), dehydroepiandrosterone (**2**), ethioallocholane (**9**), 3-deoxydehydroepiandrosterone (**10**) and 5α-androstan-3-one (**11**), the solid residues after biocatalysis were redissolved in chloroform containing 30 mM cholesterol as internal standard. In case of substrates etiadienic acid ethyl ester (**8**) samples were dissolved in pure chloroform. Samples were analyzed on a GC2010 gas chromatograph (Duisburg, Shimadzu) equipped with an OPTIMA 17ms column (Macherey-Nagel, Düren, Germany) with a linear gradient starting at 250 °C and heating with 10 °C min^−1^ until 300 °C, with the exception of substrate **9** were the gradient started at 230 °C. Injector and detector temperature were set to 350 and 300 °C respectively. Substrates and products were detected by flame ionization detector (FID). Substrates, pregnenolone (**1**), dehydroepiandrosterone (**2**), etiadienic acid ethyl ester (**8**), ethioallocholane (**9**), 3-deoxydehydroepiandrosterone (**10**) and 5α-androstan-3-one (**11**) were detected with a retention time of 9.69 min, 7.94 min, 9.36 min, 3.87 min, 4.97 min and 5.37 min, respectively, whereas their known products eluted at 9.27 (16α-OH-**1**), 10.18 min (16α-OH-**2**), 6.14 min (16α-OH-**10**) and 8.24 min (15α-OH-**11**), respectively. In case of substrate finasteride (**7**) the solid residue was re-dissolved in chloroform and samples were analyzed on a GC2010 equipped with a Supreme 5ms column (CS Chromatographie Service, Langerwehe, Germany). For separation, an isothermal temperature program at 300 °C was applied for 20 min. Injector and detector temperature were set to 300 °C. Substrate **7** was eluted with a retention time of 5.22 min.

The dried residues of bioconversions with progesterone (**3**), androstendione (**4**), testosterone (**5**) and nandrolone (**6**) were dissolved in acetonitrile: water (60:40) and injected on an ultra-fast liquid chromatograph (UFLC, Prominence, Shimadzu) equipped with a Nucleosil 100-5 C18, 250 × 4,5 mm column at 50 °C. Acetonitrile: water (60:40) was used as mobile phase with a flow rate of 1.2 mL min^−1^. Detection of substrates and their respective products was performed by UV absorbance at 243 (**5** and 16α-OH-**5**) and 242 nm (**3**, **4**, **6**, 16α-OH-**3**, 16α-OH-**4** and 16α-OH-**6**). The substrates were eluted at 5.93 min (**5**), 10.45 min (**3**), 6.35 min (**4**) and 5.39 min (**6**), whereas the main products were detected at 3.23 min (16α-OH-**5**), 4.35 min (16α-OH-**3**), 5.60 min (21-OH-**3**), 3.87 min (16α-OH-**4A**) and 3.26 min (16α-OH-**6A**). In all cases the conversions were calculated based on substrate consumption.

### GC-MS and NMR analyses

Preliminary product identification was performed with a gas chromatograph - mass spectrometer (GC-MS-QP2010S, Shimadzu, Germany) equipped with an OPTIMA 17ms column (Macherey-Nagel, Germany) using a linear temperature gradient starting at 250 °C and heating with 10 °C min^−1^ to 300 °C. Injector, interface and ion source temperature were set to 300, 300 and 200 °C respectively. Structure elucidation of formed products was performed by ^1^H-, ^13^C-, COSY, HSQC, DEPT and NOESY NMR analysis on Bruker AV400, AV500 or AV600 instruments using deuterated chloroform or DMSO as solvent with TMS as internal standard. Chemical shifts (δ) are given in ppm and coupling constant (*J*) in Hz.

### MD simulations

Molecular dynamics simulations were performed using Yasara with algorithms that were recently described in detail elsewhere.^[28]^ Periodic boundary conditions were applied. Long range electrostatics, beyond 7.86 Å, were calculated through the particle mesh Ewald method using 4th degree spline functions. The time-step was 2.0 fs, with the non-bonded interactions updated every 2 time-steps. The force-field was Yamber3, which is an Amber99 derivative which was specifically parameterized for structural accuracy.^[29]^ The compound I structure was generated and its atomic point charges assigned as described by the Pleiss group.^[30]^ Point charges on the steroid substrates were generated with AM1-BCC, which gives similar accuracy as RESP at a much lower computational cost.^[31]^

Prior to the MD simulations, an energy minimization (described previously^[32]^) was used to remove subtle steric clashes. For each of the modeled enzyme-substrate complexes, three independent MD simulations were carried out. These simulations were started with different initial atom velocities assigned via a random seed number.^[33]^ The distribution of these atom velocities always obeyed a Boltzmann distribution. In the first 30 ps of the MD simulation, the temperature was gradually increased from 5 to 298 K. After that, the simulation was allowed to equilibrate for 1970 ps. The subsequent production phase was 20 ns. From the latter phase, all the reported results were collected. Snapshots were saved every 50 ps.

During the production phase of the MD simulations, geometric information was recorded to quantify to which extent hydrogen atoms of the substrate were in a suitable orientation to be attacked by the oxygen atom of compound I. These geometries were recorded every 1 ps, on the fly. As suggested by the Bruice group, near attack conformations (NACs) were defined as having interatomic distances of less than the Van der Waals contact distance and angles between the reacting atoms within 20º of those in the quantum mechanically modeled transition state (Figure S8).^[34,35]^ A Yasara script that automatically performs the MD simulations, the recording of these NACs, and analysis of the resulting data is available upon request.

### Docking

A challenge with the flexible P450 class of enzymes is that substrate binding often requires small backbone changes. Therefore, docking was carried out essentially as previously performed by the Reetz group for modeling steroid binding and conversion by P450 BM3 variants.^[10]^ First the F92A mutation was introduced into the four experimentally determined CYP154C5 structures that have a steroid substrate bound.^[11]^ Subsequently 12 ns MD simulations were carried out, to sample the possible backbone changes around the active site. Progesterone was docked to snapshots of these MD simulations (1 snapshot was used per ns of MD simulation). The docking was performed using Autodock4^[36]^ with 4995 docking runs for each snapshot and 25000 energy evaluations per docking run. Substrate orientations with an unrealistic binding orientation were avoided by eliminating all poses of which the predicted binding energy fell outside the 90% confidence interval of the substrate orientation with the best binding energy, as described earlier.^[37]^ The same protocol was used for docking steroid **10** as well as 16α- or 4β-hydroxylated **10** into the active site of CYP154C5 but using the wild-type structures as starting points.

### Protein crystallography

Co-crystallization experiments for wild-type CYP154C5 with 5α-androstan-3-one (**11**) were set up using a HoneyBee 961 pipetting robot (Digilab Genomic Solutions, Hopkinton, U.S.A.) mixing 200 nL of protein solution (containing 40 mg/mL of enzyme and 2 mM of ligand) and 200 nL of reservoir solution, and monitored with a Rock Imager 1000 automated microscope (Formulatrix, Bedford, U.S.A.). As crystallization trails for wild-type CYP154C5 based on literature conditions^[11]^ did not directly yield crystals, sparse matrix screens were applied to identify 0.2 M ammonium sulfate, 0.1 M BisTris pH 6.5 and 25% (w/v) PEG3350 as suitable precipitant mixture. Brownish, triangle-shaped crystals were cryoprotected by soaking in reservoir solution containing 10% (v/v) 2,3-(*R*,*R*)-butanediol prior to freezing in liquid nitrogen. 3600 images with an oscillation angle of 0.1° were collected at beamline X06SA (PXI) at the Swiss Light Source (Paul Scherrer Institut, Villigen, Switzerland) on an EIGER 16M X detector.

### Structure determination

Data sets were processed using DIALS^[38]^, POINTLESS^[39]^ and AIMLESS^[40]^ of the CCP4 suite^[41]^ applying a resolution cut-off of 2 Å yielding a CC1/2 higher than 0.5 in the lowest resolution shell. Initial maps were calculated by Fourier synthesis using phenix.refine^[42]^ and the atomic coordinates of CYP154C co-crystalized with testosterone (PDB: 4j6d). The structure was further refined by alternating rounds of manual adjustment in COOT^[43]^ and computer-driven refinement with phenix.refine, including TLS refinement. Geometry restraints for 5α-androstan-3-one were calculated using the Grade Web Server (http://grade.globalphasing.org; Global Phasing Inc.). Data processing and refinement statistics are listed in Supplementary Table S3. Diffraction data and coordinates were deposited in the Protein Data Bank^[44]^ (PDB: 6TO2).

## Abbreviations

CYP: cytochrome P450 monooxygenase
ETC: electron transfer component
MD: molecular dynamics
NAC: near attack conformation
OD: optical density
PdR: putidaredoxin reductase
Pdx: putidaredoxin
TON: turnover number

## Acknowledgements

We thank Suzanne Flaton (HIMS, University of Amsterdam) for assistance with preparative-scale reactions. We also thank Ines Bachmann-Remy (ITMC, RWTH Aachen University) and Jan Meine Ernsting (HIMS, University of Amsterdam) for technical assistance with NMR measurements. We further acknowledge the Paul Scherrer Institute, Villigen, Switzerland for provision of synchrotron radiation beamtime at beamline X06SA of the SLS and would like to thank Dr. Takashi Tomizaki for assistance.

This project was financially supported by the German Research Foundation (DFG) within the national Excellence Initiative funding scheme to promote science and research at German universities. Additionally, financial support of JARB and TK by the German Research Foundation via the Research Training Group PROCOMPAS [GRK 2223] is gratefully acknowledged.

## Author Contributions

AS, HJW and DBJ designed the work; PB, HJW, BN, AV, PS and JARB performed experiments; PB, HJW, PS, TK, JARB and AS analyzed the data; PB, HJW, TK and AS wrote the manuscript; AS, WB and DBJ revised it critically. All authors approved the final version for publication.

