## Supporting information for "CYP154C5 Regioselectivity in Steroid Hydroxylation Explored by Substrate and Protein Engineering"

### 1. Mutagenesis

**Table S1.** Applied QuikChange® primers for the generation of CYP154C5 mutants. The introduced alanine codon is underlined.

| Primer | Sequence (5' to 3') |
|---|---|
| M84A fwd | GCCGCTGATCGGCGCGATCGACGTGGAC |
| M84A rev | GTCCACGTCGATCGCGCCGATCAGCGGC |
| F92A fwd | GGACCGCTCGATGGCCACCGTGGACGGC |
| F92A rev | GCCGTCCACGGTGGCCATCGAGCGGTCC |
| Q239A fwd | TGATCGGCAATCTCGCGGCGCTCGTCGCC |
| Q239A rev | GCGACGAGCGCCGCGAGATTGCCGATCAG |
| Q398A fwd | CCCGTCCTCACCGCGAACGACCTGTCCCAC |
| Q398A rev | GTGGGACAGGTCGTTCGCGGTGAGGACGGG |

### 2. Protein purification

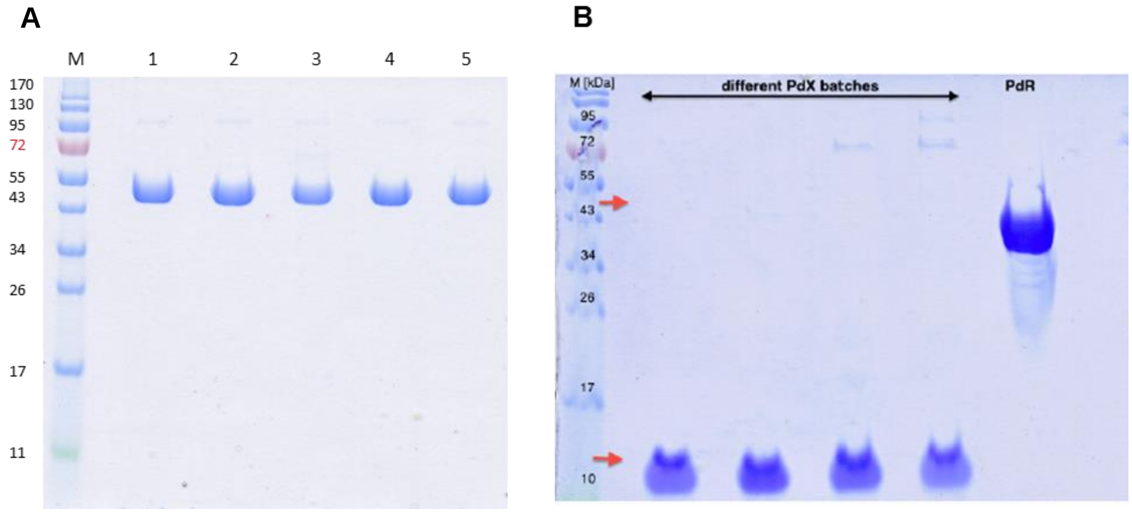


**Figure S1.** SDS-PAGE of purified proteins. **M** represents the Marker; Proteins shown correspond to **A**) 1: CYP154C5 (45.3 kDa); 2: CYP154C5 M84A; 3: CYP154C5 Q398A; 4: CYP154C5 Q239A; 5: CYP154C5 F92A and **B**) four different Pdx (11.5 kDa) batches; PdR (45.5 kDa).

### 3. Biochemical characterization of CYP154C5 mutants

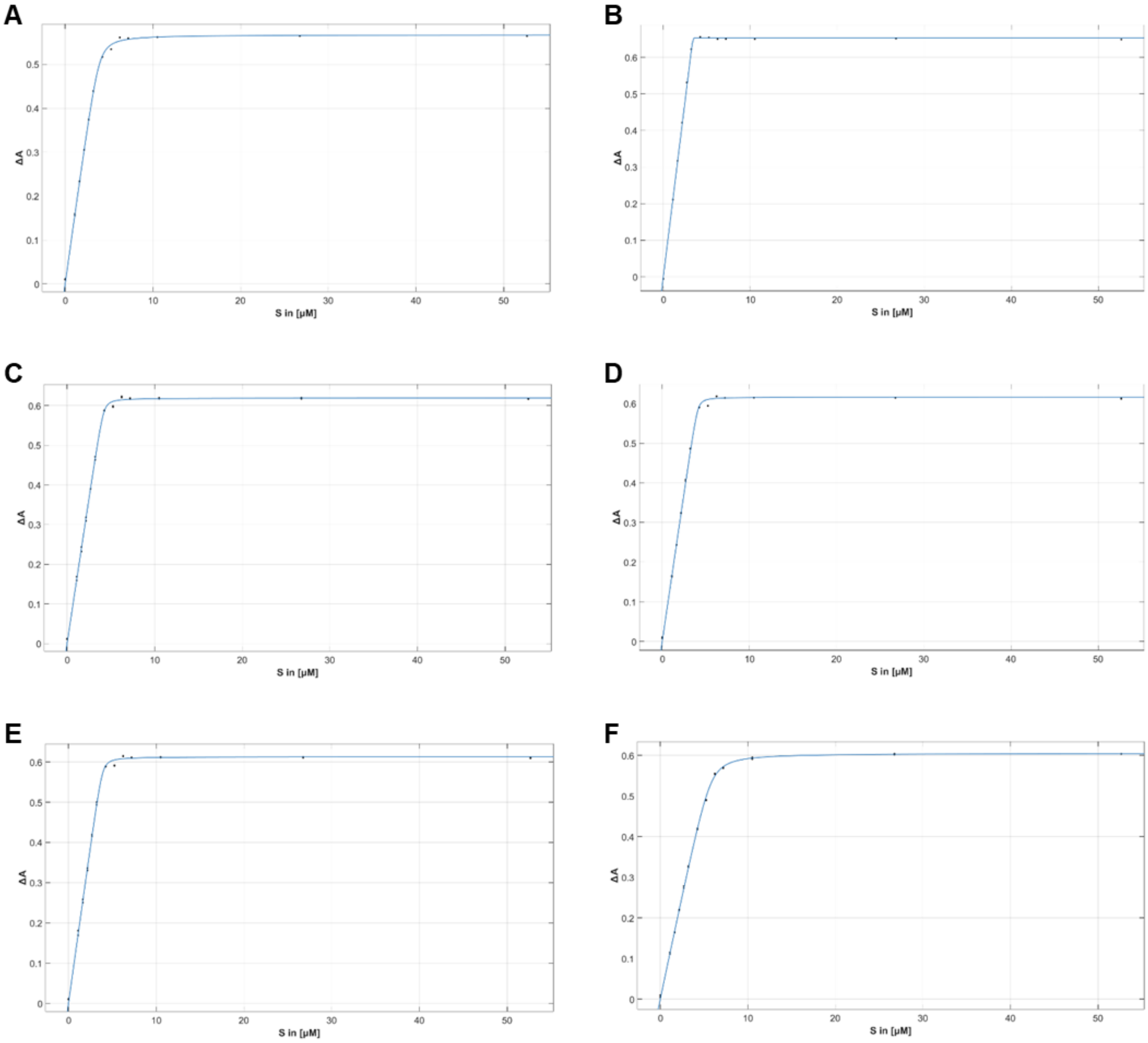


**Figure S2.** Substrate-binding titrations for CYP154C5 wild type using steroid substrates **A**: pregnenolone (**1**), **B**: dehydroepiandrosterone (**2**), **C**: progesterone (**3**), **D**: androstenedione (**4**), **E**: testosterone (**5**) and **F**: nandrolone (**6**). ΔA was plotted against the applied steroid concentration and the resulting data was fitted using the tight binding equation.

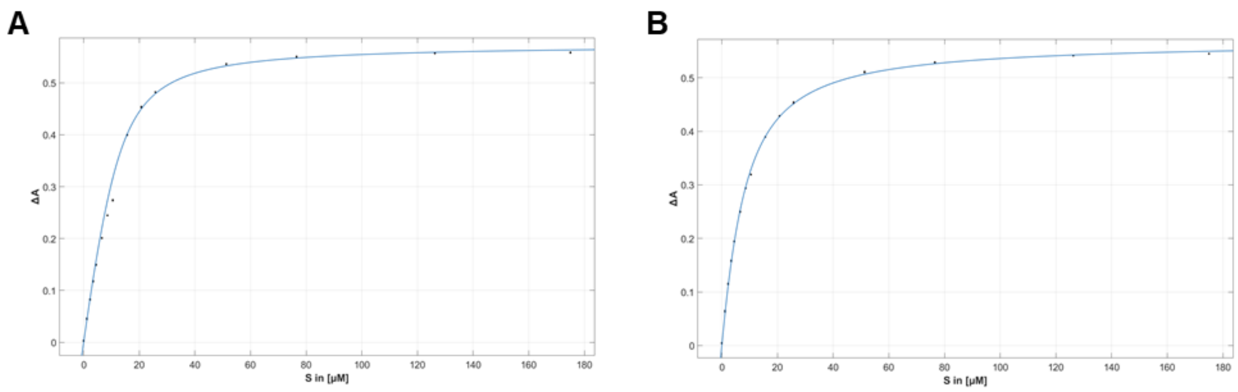


**Figure S3.** Substrate-binding titrations for CYP154C5 M84A using steroid substrates **A**: androstenedione (**4**) and **B**: testosterone (**5**). ΔA was plotted against the applied steroid concentration and the resulting data was fitted using the tight binding equation.

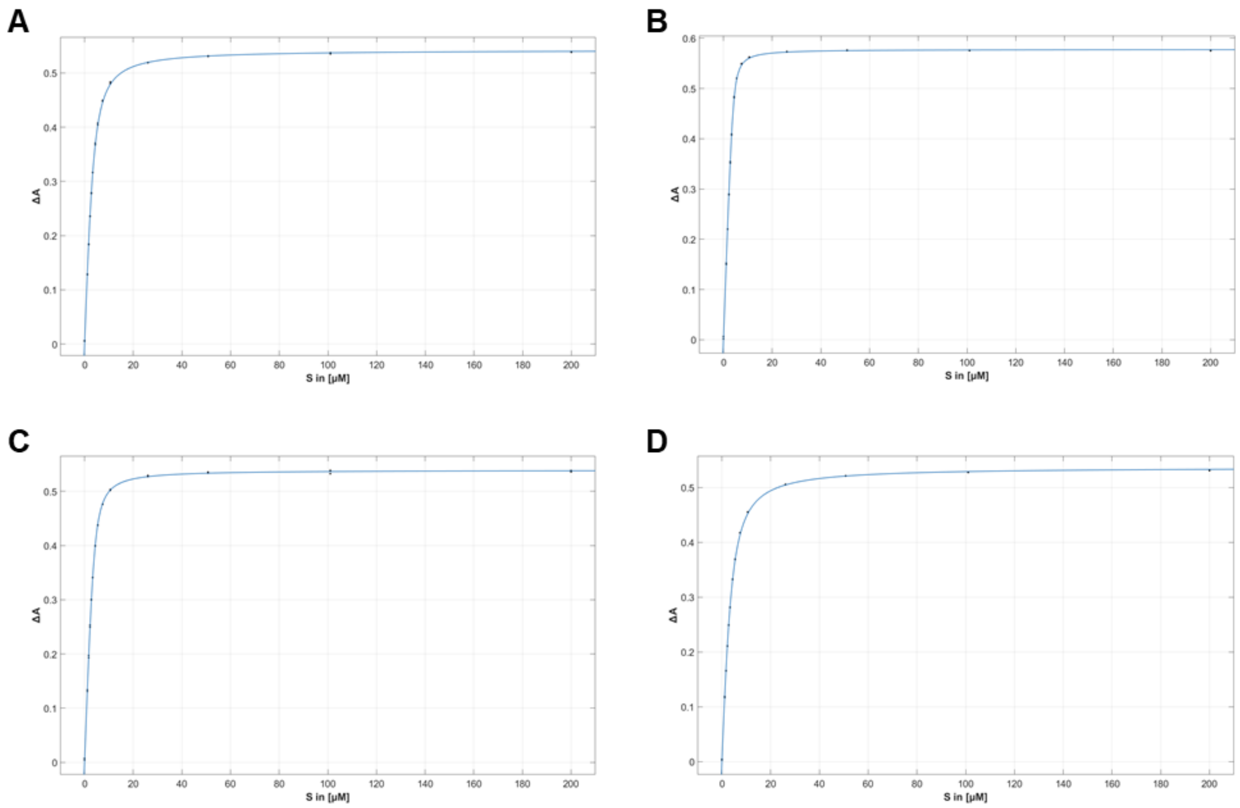


**Figure S4.** Substrate-binding titrations for CYP154C5 F92A using steroid substrates **A**: dehydroepiandrosterone (**2**), **B**: progesterone (**3**), **C**: androstenedione (**4**) and **D**: testosterone (**5**). ΔA was plotted against the applied steroid concentration and the resulting data was fitted using the tight binding equation.

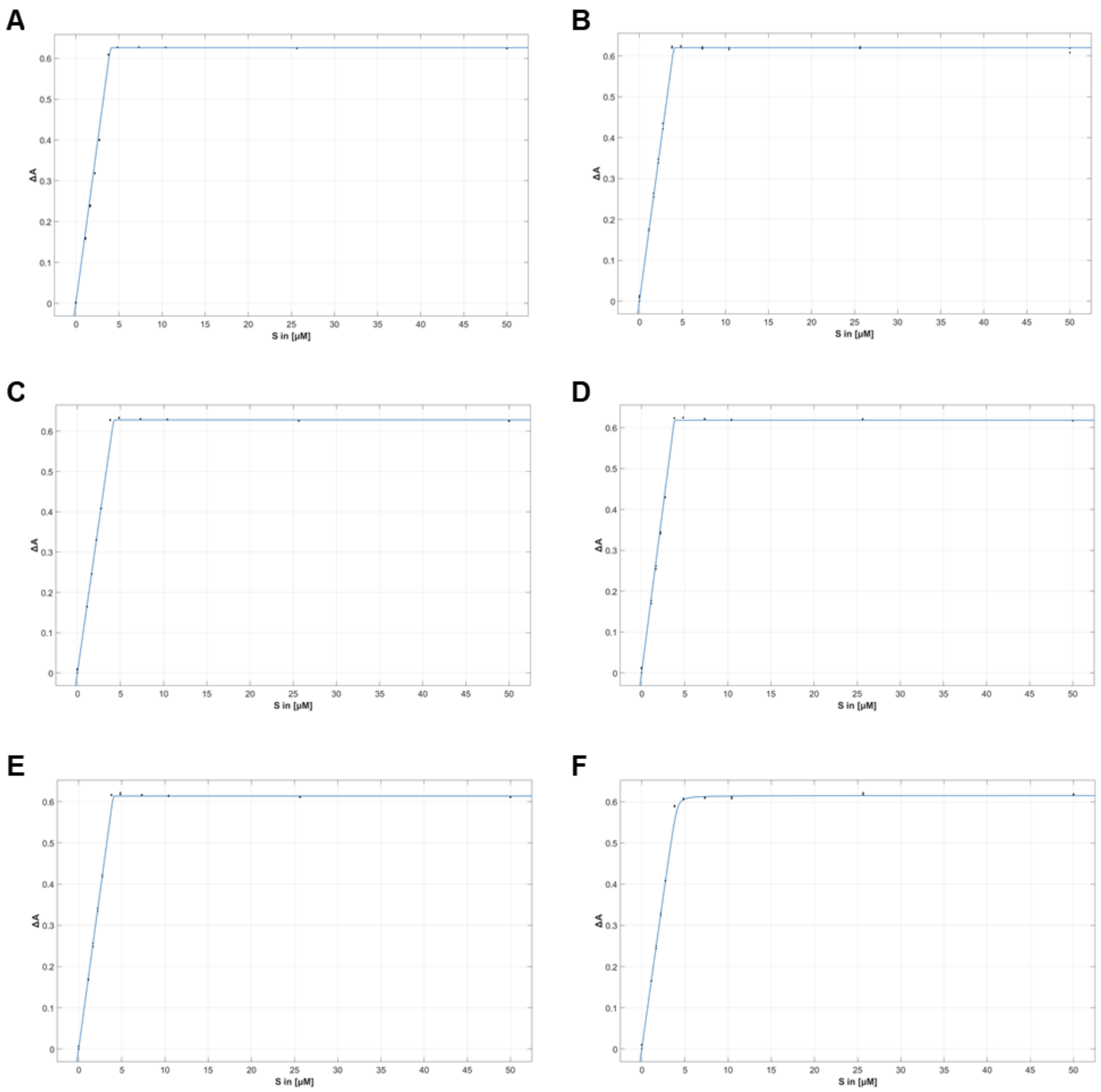


**Figure S5.** Substrate-binding titrations for CYP154C5 Q239A using steroid substrates **A**: pregnenolone (**1**), **B**: dehydroepiandrosterone (**2**), **C**: progesterone (**3**), **D**: androstenedione (**4**), **E**: testosterone (**5**) and **F**: nandrolone (**6**). ΔA was plotted against the applied steroid concentration and the resulting data was fitted using the tight binding equation.

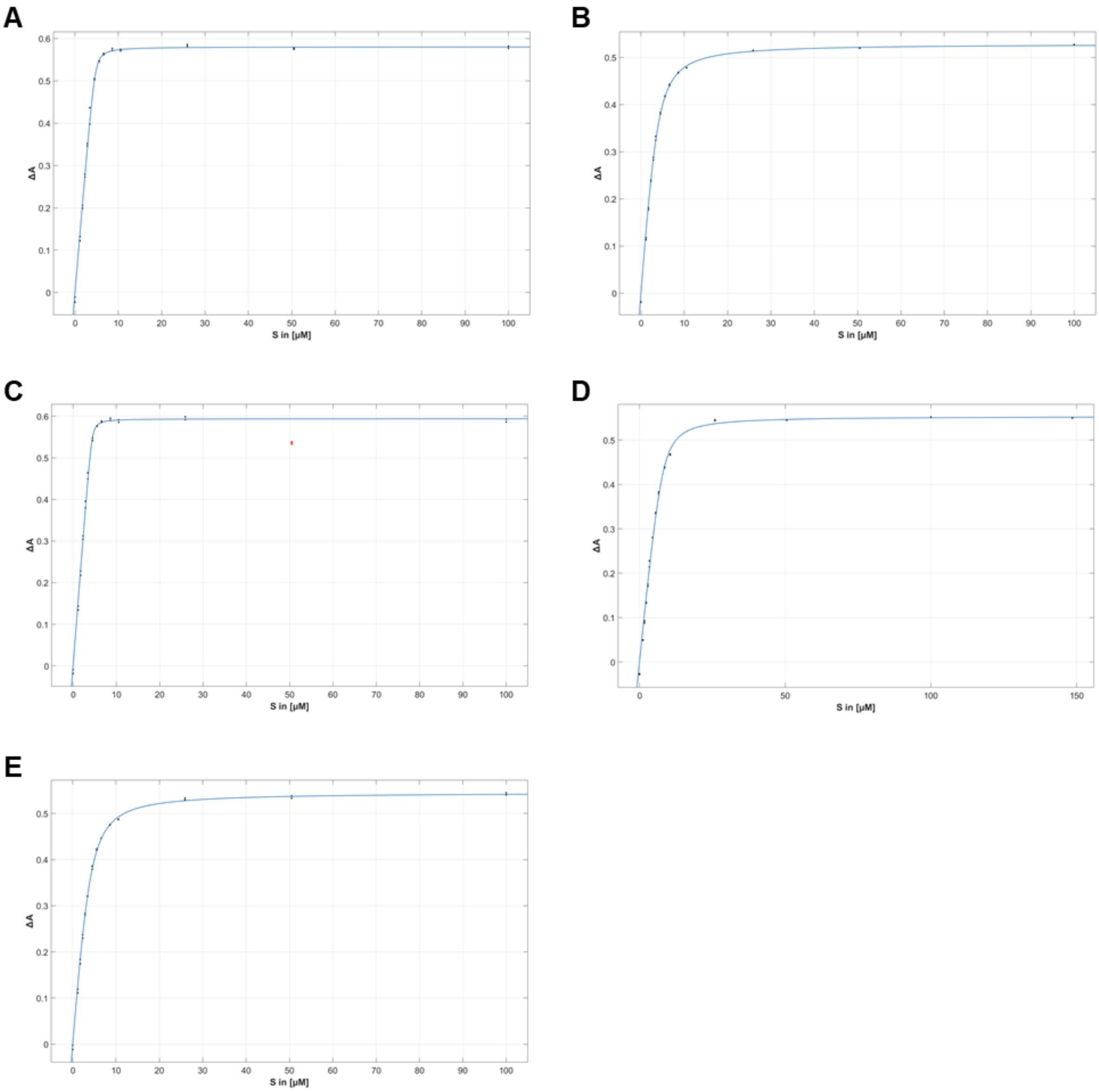


**Figure S6.** Substrate-binding titrations for CYP154C5 Q398A using steroid substrates **A**: pregnenolone (**1**), **B**: dehydroepiandrosterone (**2**), **C**: progesterone (**3**), **D**: androstenedione (**4**) and **E**: testosterone (**5**). ΔA was plotted against the applied steroid concentration and the resulting data was fitted using the tight binding equation. Red dots represent data points that were not included in the fitting.

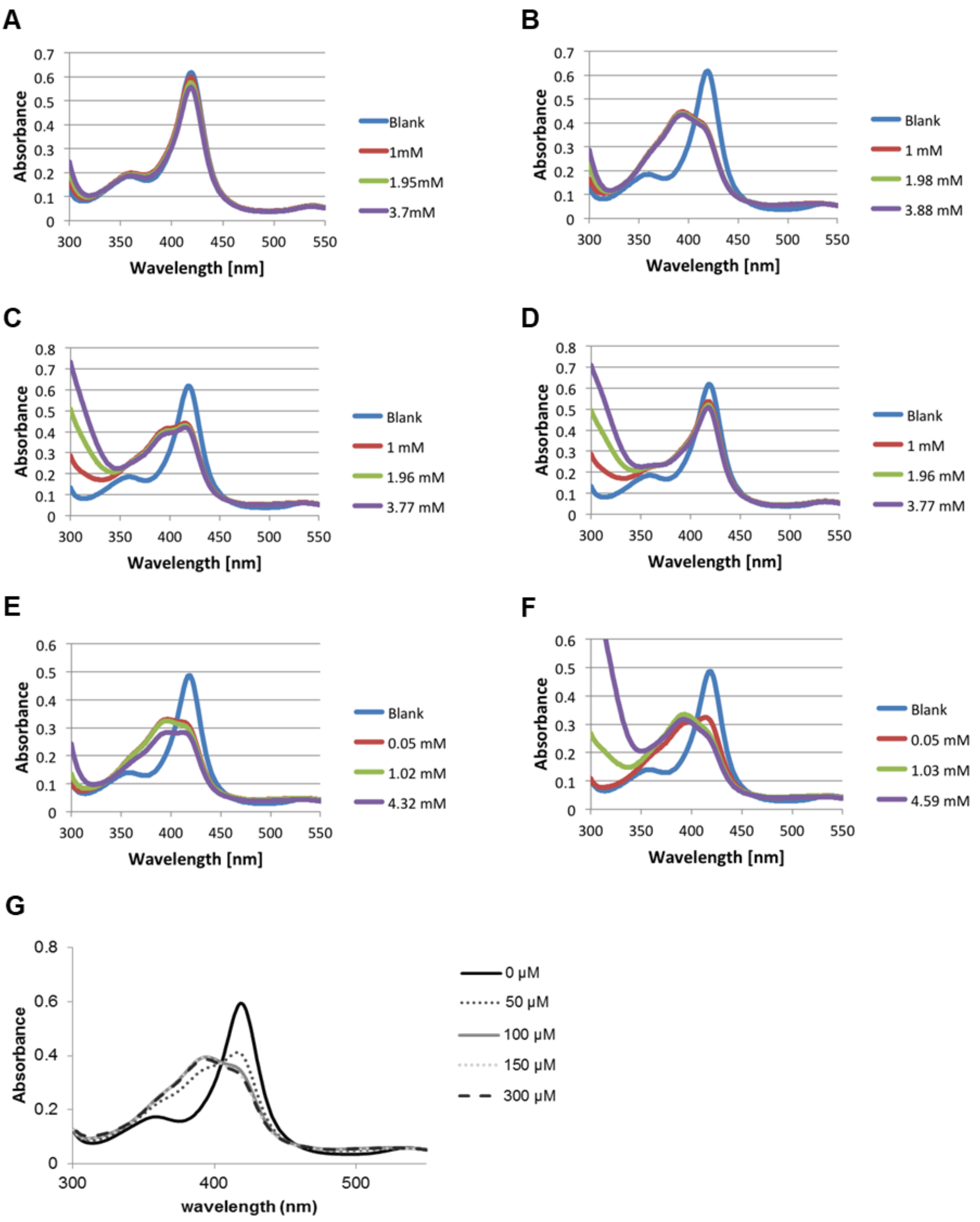


**Figure S7.** Substrate titrations for $K_D$ determination. The following enzyme-substrate combinations resulted only in a partial or no spectral shift upon substrate addition: (**A**) CYP154C5 M84A with **1**; (**B**) CYP154C5 M84A with **2**; (**C**) CYP154C5 M84A with **3**; (**D**) CYP154C5 M84A with **6**; (**E**) CYP154C5 F92A with **1**; (**F**) CYP154C5 F92A with **6** and (**G**) CYP154C5 WT with **9**.

**Table S2.** Obtained conversion values for purified CYP154C5 wild type and mutants in the transformation of steroids **1-6** at 30°C after 8 h reaction time. Measurements were performed in duplicate.

| | **Conversion (%)** | | | | |
|---|---|---|---|---|---|
| **Substrate** | **CYP154C5** | **CYP154C5 M84A** | **CYP154C5 F92A** | **CYP154C5 Q239A** | **CYP154C5 Q398A** |
| Pregnenolone (**1**) | 100 ± 0 | 8 ± 5 | 27 ± 1 | 99 ± 0 | 40 ± 2 |
| Dehydroepi-androsterone (**2**) | 100 ± 0 | 26 ± 1 | 83 ± 12 | 100 ± 0 | 39 ± 0 |
| Progesterone (**3**) | 100 ± 0 | 67 ± 1 | 83 ± 1 | 100 ± 0 | 82 ± 1 |
| Androstene-dione (**4**) | 100 ± 0 | 26 ± 0 | 59 ± 0 | 87 ± 1 | 100 ± 0 |
| Testosterone (**5**) | 74 ± 2 | 38 ± 1 | 58 ± 1 | 100 ± 0 | 28 ± 3 |
| Nandrolone (**6**) | 69 ± 3 | –[a] | 32 ± 0 | 100 ± 0 | –[a] |

[a] No conversion observed

### 4. CYP154C5 modeling and docking

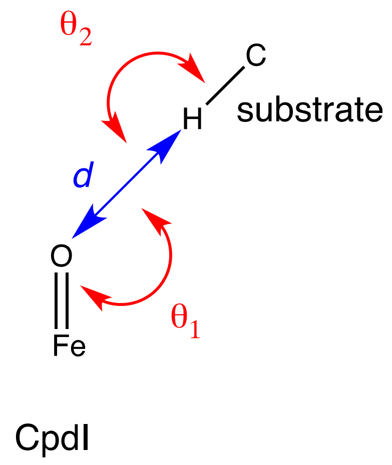


**Figure S8.** Geometric definitions of a near attack conformation for P450-catalyzed hydroxylation. Of the P450, only the iron and the reactive oxygen atom of compound I are shown while of the substrate only the attacked hydrogen and carbon atom are shown. A conformation was scored to be a NAC if it displayed simultaneously a distance $d \leq 2.72$ Å, an angle $\theta_1$ of 100-140°, and an angle $\theta_2$ of > 140°.

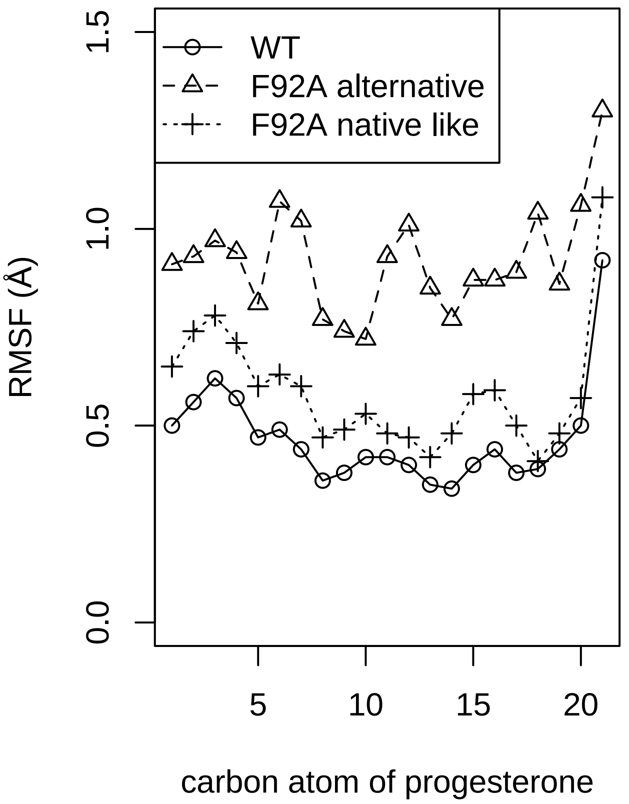


**Figure S9.** The positional flexibility of the substrate progesterone (**3**) increases due to the F92A mutation in CYP154C5. For each complex, the RMSF was calculated from three independent MD simulations of each 22 ns.

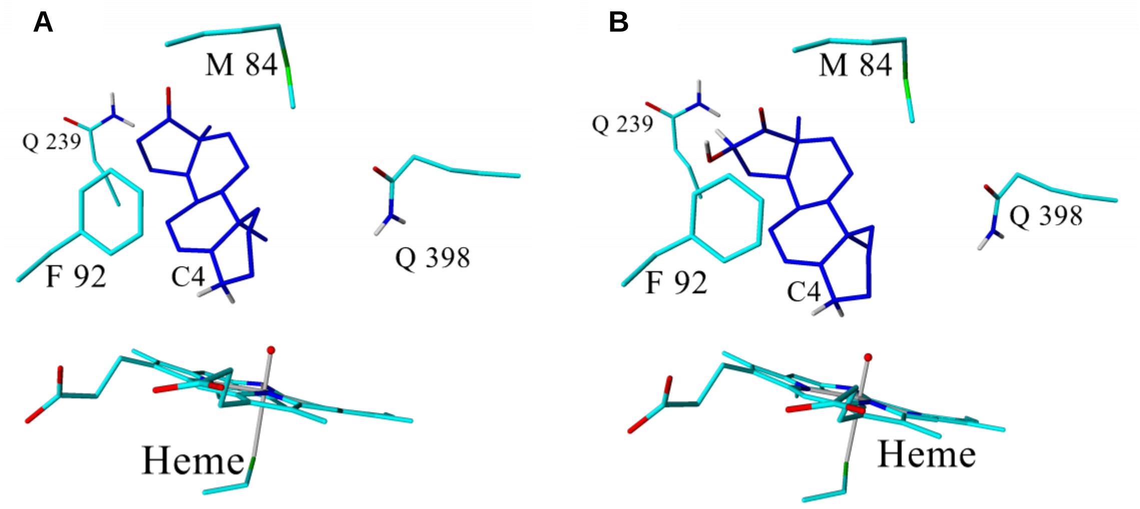

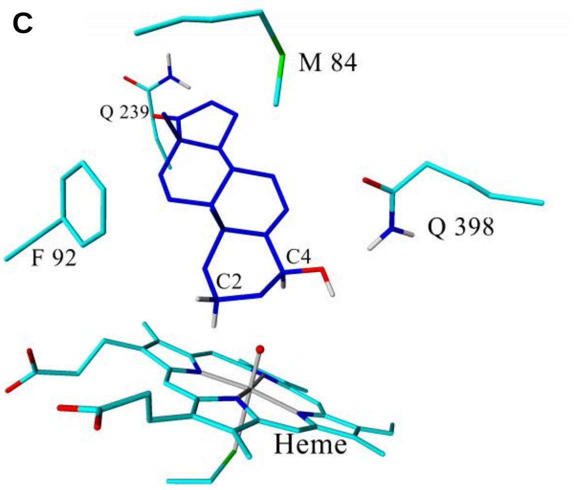


**Figure S10.** Obtained steroid orientations and potential hydroxylation sites during docking of **A**) 3-deoxydehydroepiandrosterone (**10**), **B**) 16α-hydroxy-3-deoxydehydroepiandrosterone and **C**) 4β-hydroxy-3-deoxydehydroepiandrosterone as substrates in CYP154C5 (as compound I model).

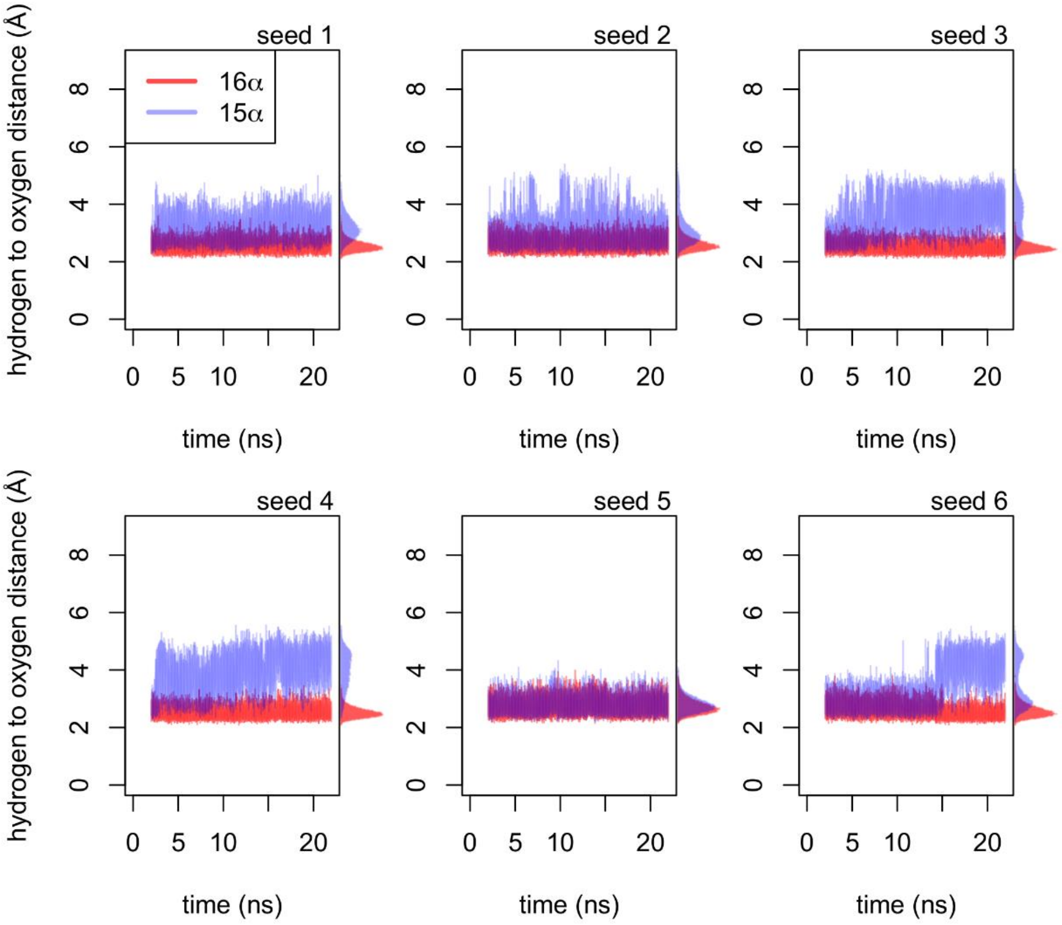

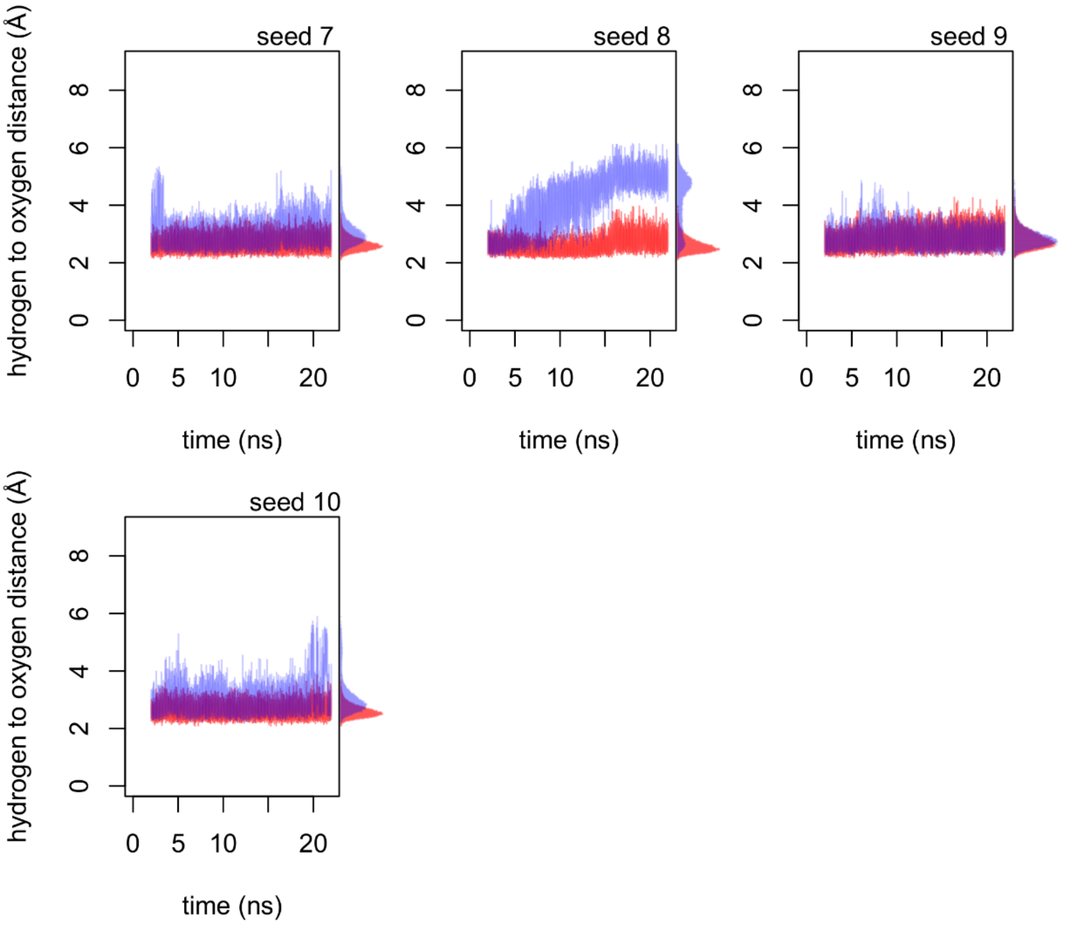


**Figure S11.** Distances of the substrate's hydrogen atoms to the oxygen of compound I during MD simulation. Since significant differences were observed within the 10 individual MD trajectories, the results of all trajectories are shown individually.

### 5. Protein crystallography

**Table S3.** Data collection and refinement statistics for CYP154C5 co-crystallized with 5α-androstan-3-one (**11**). Values in parentheses are for the highest resolution shell.

| **Data collection** | |
|---|---|
| Wavelength | 1.000 |
| Resolution range | 46.6 - 2.0 (2.072 - 2.0) |
| Space group | R 3 H |
| Unit cell | 103.411 103.411 218.21 90 90 120 |
| Total reflections | 618601 (58073) |
| Unique reflections | 58736 (5859) |
| Multiplicity | 10.5 (9.9) |
| Completeness (%) | 99.84 (99.62) |
| Mean I/sigma(I) | 17.11 (0.96) |
| Wilson B-factor | 31.29 |
| R-merge | 0.3117 (1.848) |
| R-meas | 0.3274 (1.949) |

| | |
|---|---|
| R-pim | 0.09959 (0.614) |
| CC1/2 | 0.976 (0.496) |
| CC* | 0.994 (0.814) |
| **Model refinement** | |
| Reflections used in refinement | 58678 (5842) |
| Reflections used for R-free | 2989 (303) |
| R-work | 0.2073 (0.2963) |
| R-free | 0.2479 (0.3520) |
| CC(work) | 0.924 (0.716) |
| CC(free) | 0.885 (0.623) |
| Number of non-hydrogen atoms | 6917 |
| macromolecules | 6286 |
| ligands | 128 |
| solvent | 503 |
| Protein residues | 810 |
| RMS(bonds) | 0.009 |
| RMS(angles) | 0.85 |
| Ramachandran favored (%) | 97.63 |
| Ramachandran allowed (%) | 2.73 |
| Ramachandran outliers (%) | 0.00 |
| Rotamer outliers (%) | 0.30 |
| Clashscore | 2.19 |
| Average B-factor | 38.27 |
| macromolecules | 38.35 |
| ligands | 28.54 |
| solvent | 39.76 |
| Number of TLS groups | 15 |

### 6. Product identification

#### a. Conversion of progesterone (3) by CYP154C5 F92A

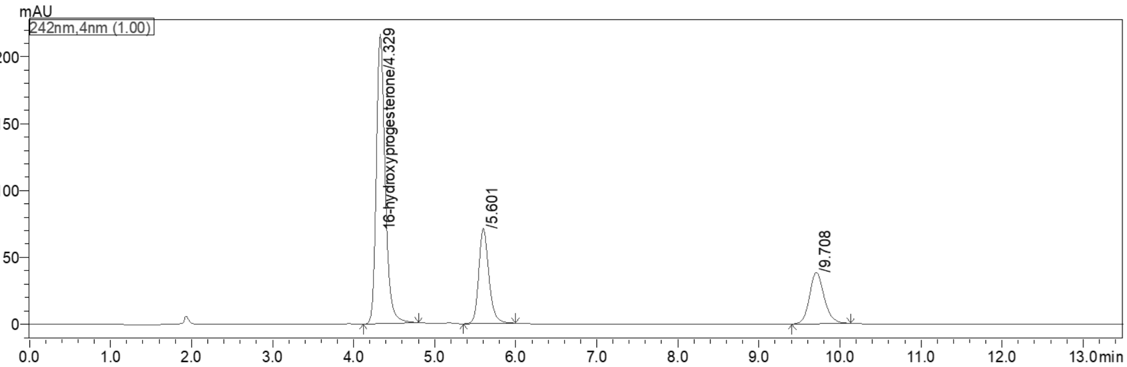


**Figure S12.** HPLC chromatogram of progesterone (**3**) conversion after 6 h reaction time catalyzed by CYP154C5 F92A. The peaks at 9.7 min and 4.3 min correspond to the substrate progesterone and the product 16α-hydroxyprogesterone, respectively. The peak at 5.6 min represents a new product identified as 21-hydroxylated progesterone by NMR analysis.

**21-hydroxyprogesterone (11-deoxycorticosterone)** $^1$H-NMR (500 MHz, $CDCl_3$): δ 5.76 (s, 1H), 4.30 - 4.14 (m, 2H), 3.30 (s, 1H), 2.55 - 2.18 (m, 7H), 2.06 (dt, *J* = 13.6, 4.2, 1H), 1.96 (dd, *J* = 9.3, 6.4, 1H), 1.89 (ddt, *J* = 11.9, 5.8, 2.7, 1H), 1.83 - 1.54 (m, 6H), 1.21 (s, 4H), 1.10 (qt, *J* = 12.9, 6.6, 2H), 1.00 (td, *J* = 11.3, 4.0, 1H), 0.91 (t, *J* = 6.8 Hz, 1H), 0.85 (dd, *J* = 13.4, 6.2, 1H), 0.72 (s, 3H). $^{13}$C-NMR (126 MHz, $CDCl_3$): 210.2 (20-C), 199.5 (3-C), 170.8 (5-C), 124.0 (4-C), 69.43 (21-C), 59.1 (17-C), 56.1 (14-C), 53.6 (9-C), 44.7 (13-C), 38.6 (10-C), 38.4 (12-C), 35.7 (8-C), 35.6 (1-C), 33.9 (2-C), 32.7 (6-C), 31.9 (7-C), 24.5 (16-C), 22.9 (15-C), 20.9 (11-C), 17.3 (19-C), 13.5 (18-C). Molecular weight: 330 g mol$^{-1}$, GC-MS: retention time: 10.5 min, $M^+$ = 312, m/z (%): 312 (28), 269 (37), 145 (16), 135 (17), 124 (24), 105 (16), 93 (16), 91 (26), 79 (17), 77 (16), 43 (100), 41 (14).

Obtained NMR and GC-MS results of formed 11-deoxycorticosterone are consistent with previously published data.[1, 2]

### b. Conversion of ethioallocholane (9) by CYP154C5 WT

**A**

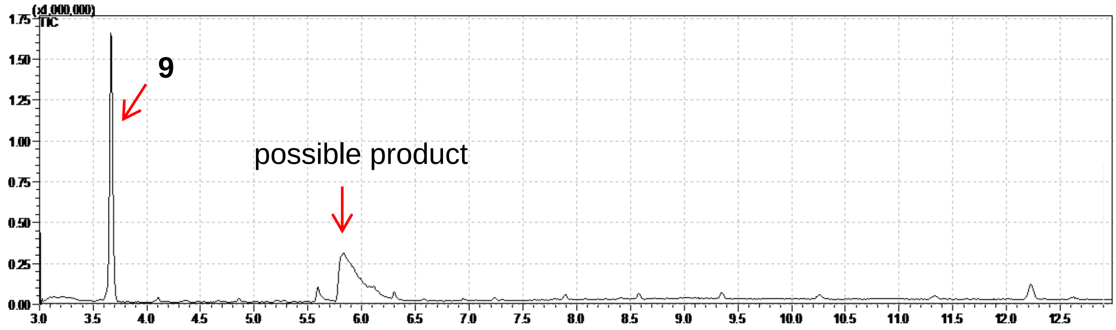


**B**

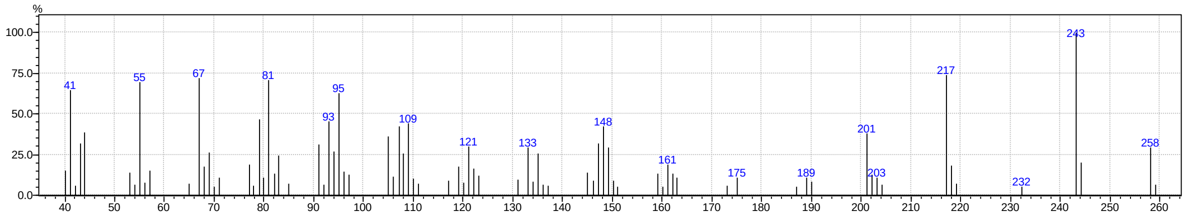


**Figure S13.** GC-MS result of ethioallocholane (**9**) conversion by CYP154C5. **A:** Chromatogram of reaction; **B:** MS of possible product peak at RT = 5.9 min with a maximum molecular ion peak of m/z = 258, equivalent to $[M]^+$-2. This likely corresponds to a hydroxylated product of **9** that undergoes water elimination during GC-MS measurement (which was already observed for other hydroxylated steroids previously).

#### c. Conversion of 3-deoxydehydroepiandrosterone (10) by CYP154C5 WT

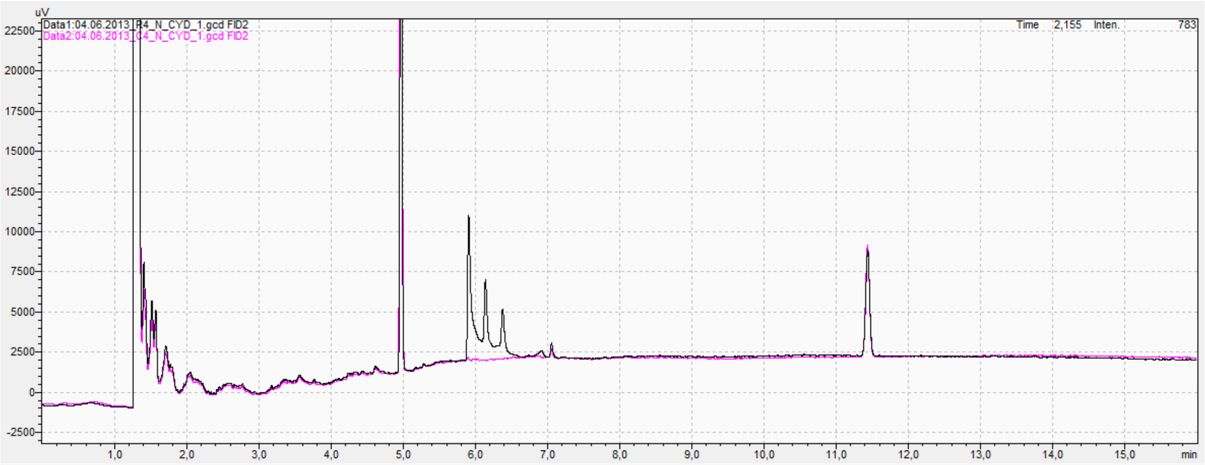


**Figure S14.** GC chromatogram (black) of 3-deoxydehydroepiandrosterone (**10**) conversion after 16 h catalysed by CYP154C5 wild type. For comparison, the GC chromatogram of the respective control reaction (magenta) lacking CYP154C5 is shown as well. The peak at 4.97 min corresponds to the substrate whereas the peaks at 5.8 to 6.6 min represent reaction products.

<u>*Main product:*</u>

**16α-hydroxy-3-deoxydehydroepiandrosterone** $^{1}$H-NMR (600 MHz, DMSO): δ 5.34 (1H, d, *J*= 5.6, 16-OH), 5.26 (1H, d, *J*=5.1, 6-H), 4.21 (1H, dd, $J_d$= 8.4, $J_d$= 5.9, 16-βH), 2.22 (1H, m, 4α-H), 2.05-1.95 (2H, m, 4β-H and 7β-H), 1.91 (1H, dt, $J_t$= 13.7, $J_d$= 8.5, 15β-H), 1.81 (1H, d, *J*= 12.7, 1α-H), 1.73-1.51 (6H, m, 2α-H, 3β-H, 8β-H, 11α-H, 12β-H and 15α-H), 1.50-1.43 (2H, m, 2β-H and 14α-H), 1.39 (1H, dq, $J_q$= 13.4, $J_d$= 4.6, 11β-H), 1.31-1.21 (2H, m, 7α-H, 12α-H), 1.14 (1H, m, 3α-H), 1.03-0.95 (5H, m, 1β-H, 9α-H and 19-Me), 0.86 (3H, m, 18-Me). $^{13}$C-NMR (150 MHz, DMSO): δ 118.9 (6-C), 70.6 (16-C), 50.6 (9-C), 48.7 (14-C), 47.2 (13-C), 39.4 (1-C), 37.6 (10-C), 32.8 (4-C), 32.1 (7-C), 31.8 (12-C), 31.3 (8-C), 30.5 (15-C), 27.9 (3-C), 22.5 (2-C), 19.8 (11-C), 19.7 (19-C), 14.3 (18-C). Molecular weight: 288 g mol$^{-1}$; GC-MS: Retention time: 6.0 min. M$^{+}$: 288; m/z (%): 288 (100), 201 (82), 216 (79), 145 (77), 41 (74), 91 (72), 105 (69), 79 (66), 55 (58), 159 (54), 121 (43).

A

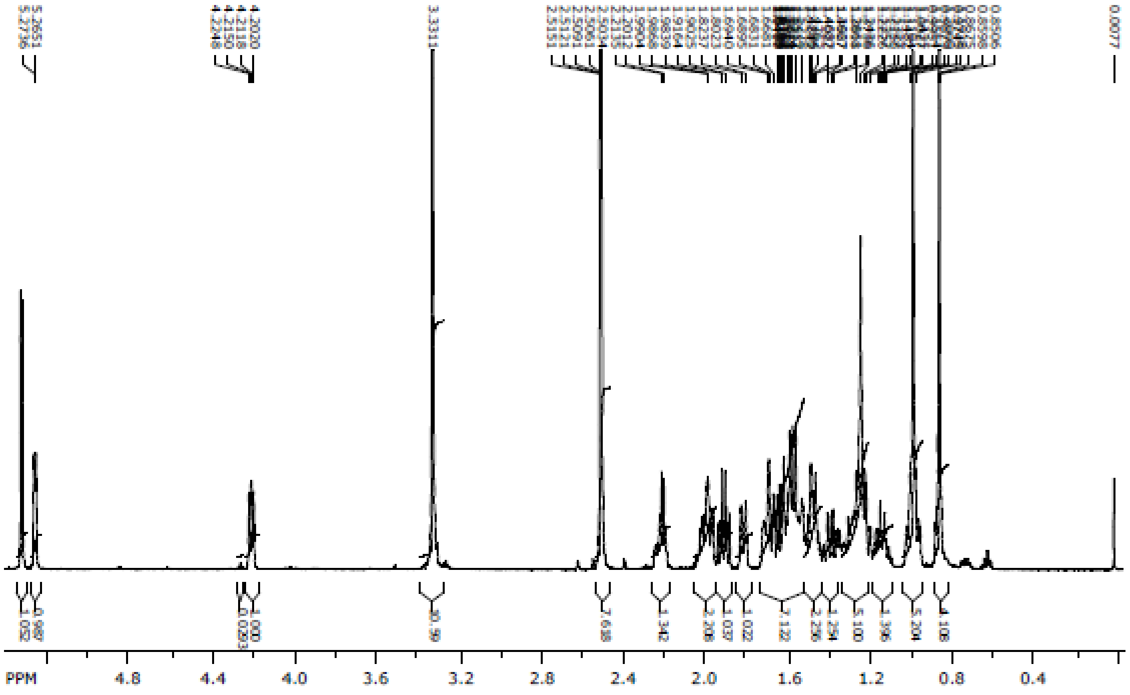

B

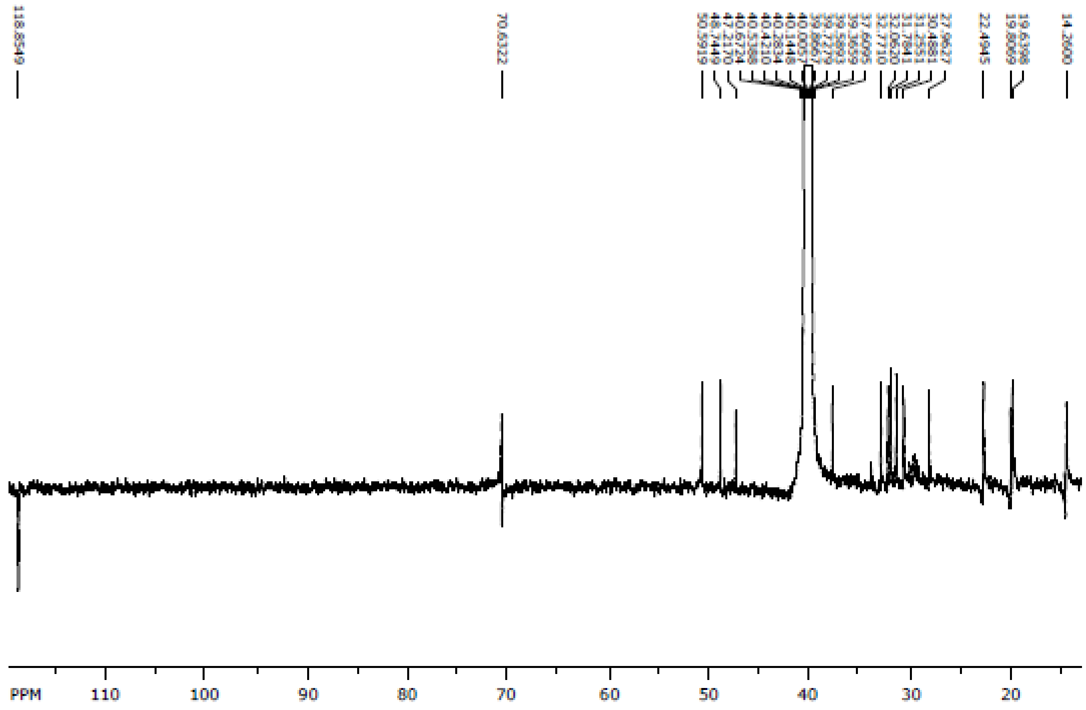

**C**

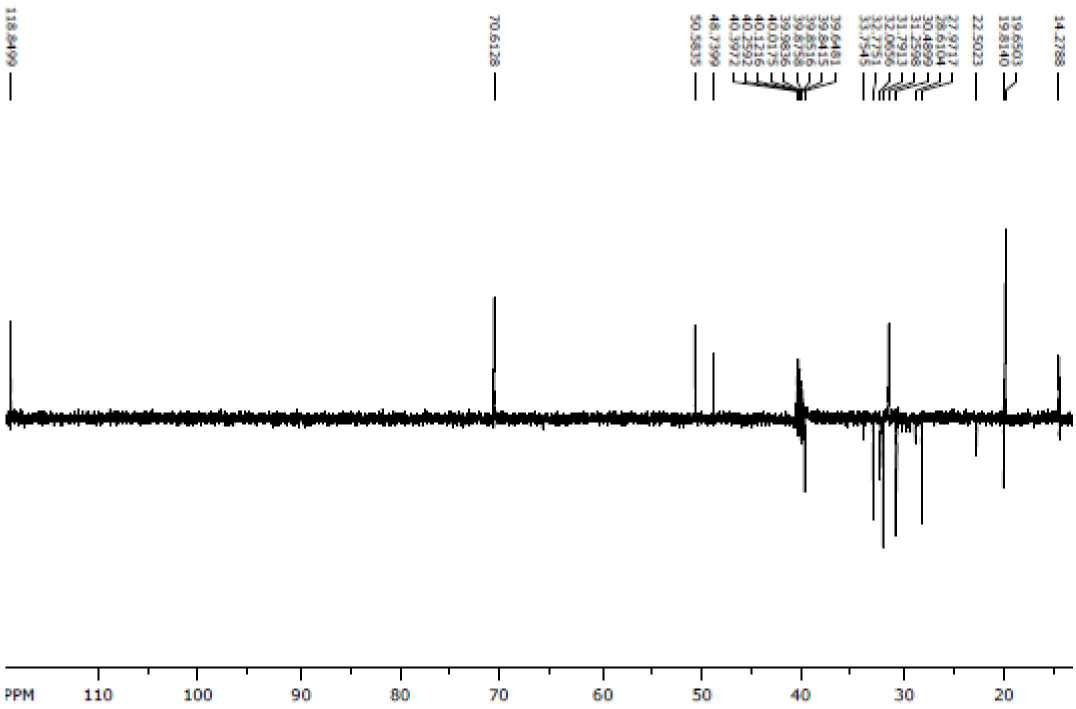
PPM
110
100
90
80
70
60
50
40
30
20

**D**

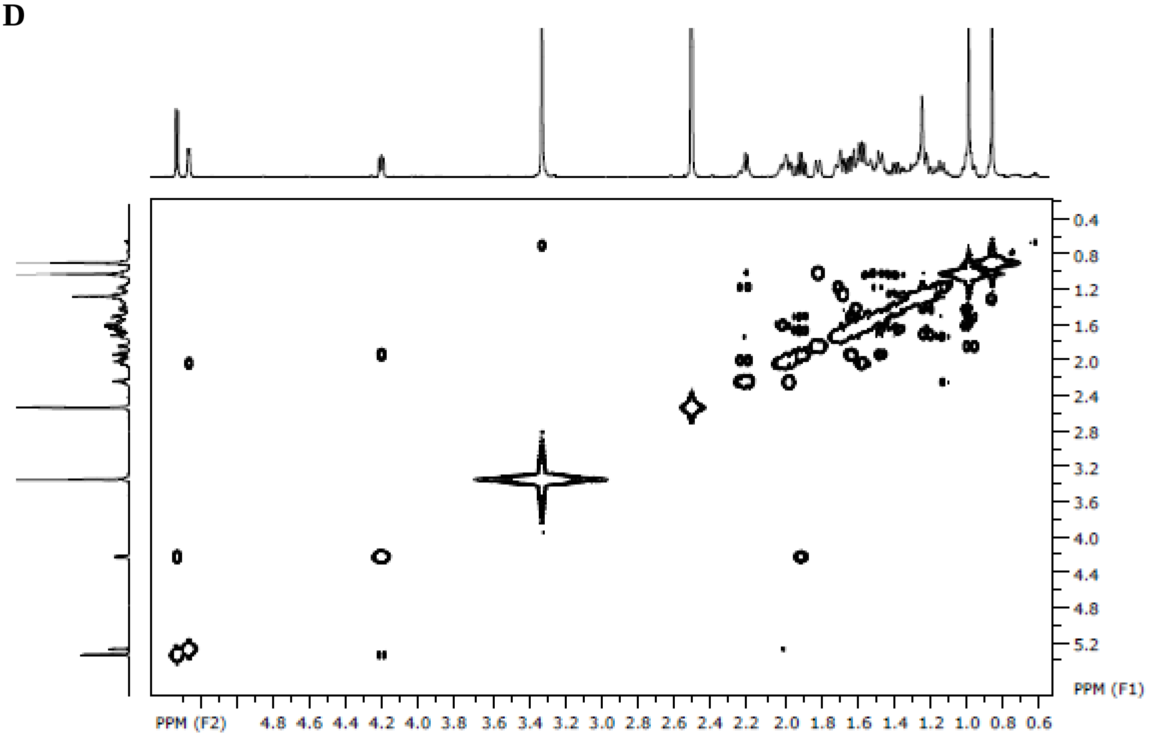
PPM (F2)
4.8 4.6 4.4 4.2 4.0 3.8 3.6 3.4 3.2 3.0 2.8 2.6 2.4 2.2 2.0 1.8 1.6 1.4 1.2 1.0 0.8 0.6
PPM (F1)
0.4 0.8 1.2 1.6 2.0 2.4 2.8 3.2 3.6 4.0 4.4 4.8 5.2

**E**

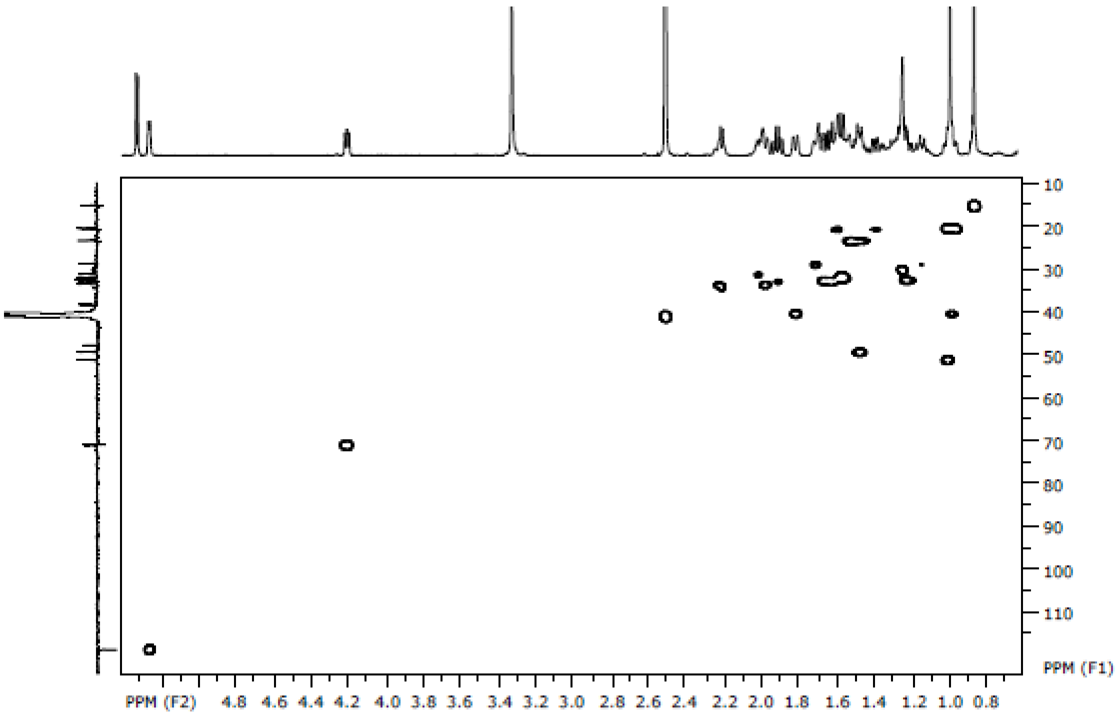


**F**

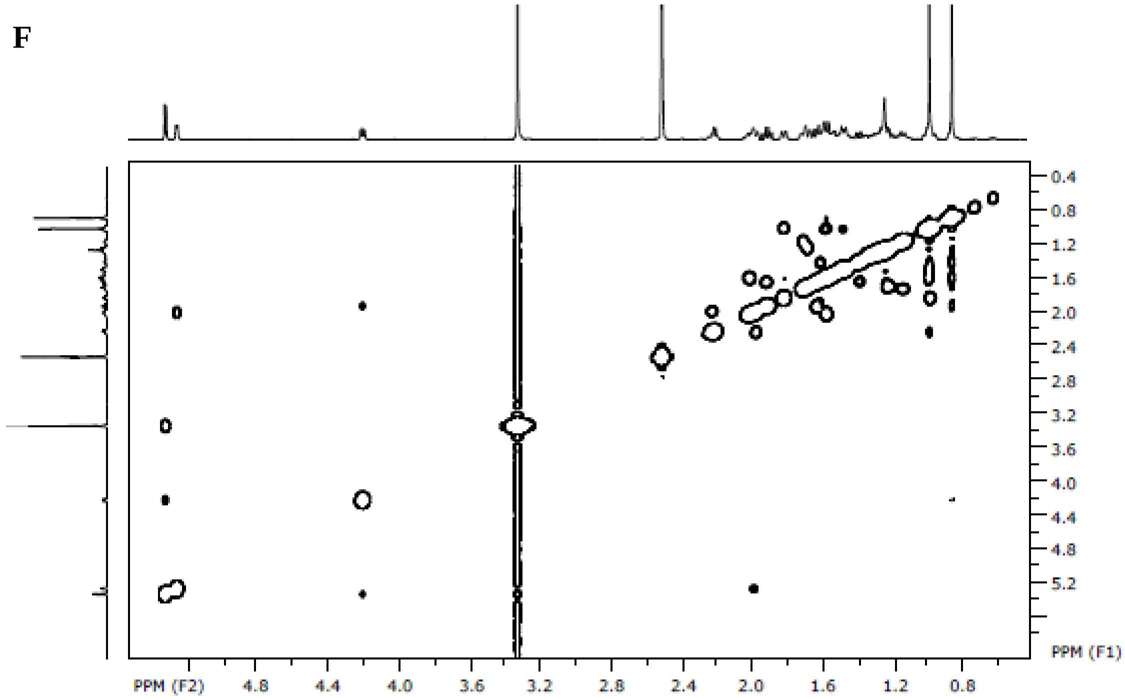


**Figure S15.** NMR spectra of 16α-hydroxy-3-deoxydehydroepiandrosterone; **A**: $^{1}$H, **B**: $^{13}$C, **C**: DEPT135, **D**: COSY, **E**: HSQC, **F**: NOESY.

*Additional 3-deoxydehydroepiandrosterone product:*

GC-MS analysis of the product gave a peak with RT = 8.7 min with a maximum molecular ion peak of m/z = 304, equivalent to $[M]^{+}$+32 (Figure S15). This result suggests a possible double hydroxylation of substrate **10**. This is consistent with the obtained $^{1}$H-NMR data showing two different hydroxylation signals at 4.42 and 4.05 ppm (Figure S16). Additionally, using HSQC-NMR analysis, these two proton signals were found to couple with carbon signals at 71.7 and 66.5 ppm corresponding to <u>C</u>H-OH (Figure S8). Unfortunately, further structure elucidation was not possible due to the low amounts of purified product and, hence, a low resolution of the NMR spectra.

**A**

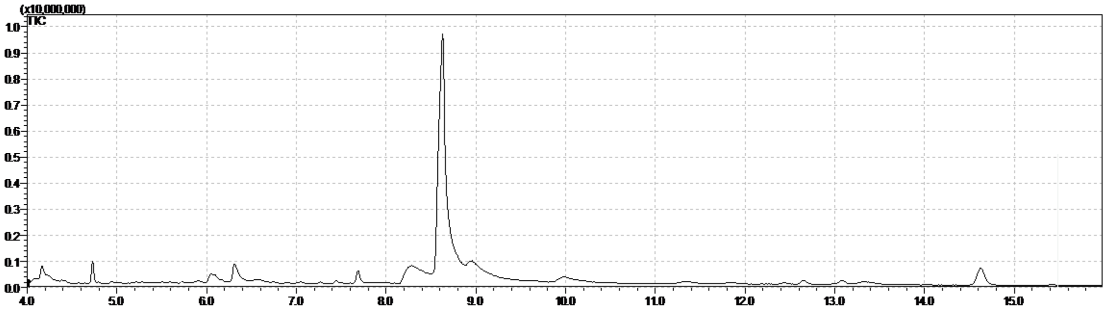


**B**

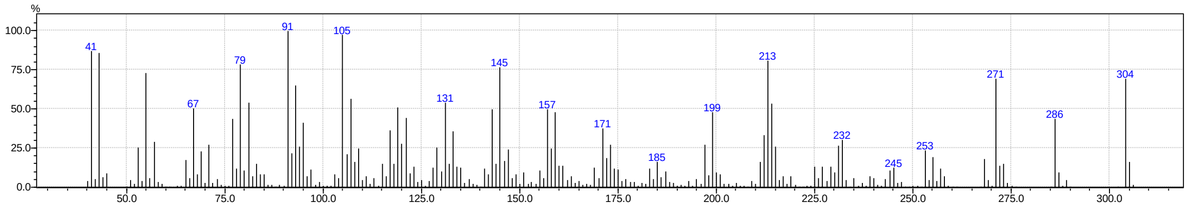


**Figure S16.** GC-MS result of the potential dihydroxylated product in the conversion of **10. A:** Chromatogram of the product. **B:** MS of the product peak at RT = 8.7 min.

**A**

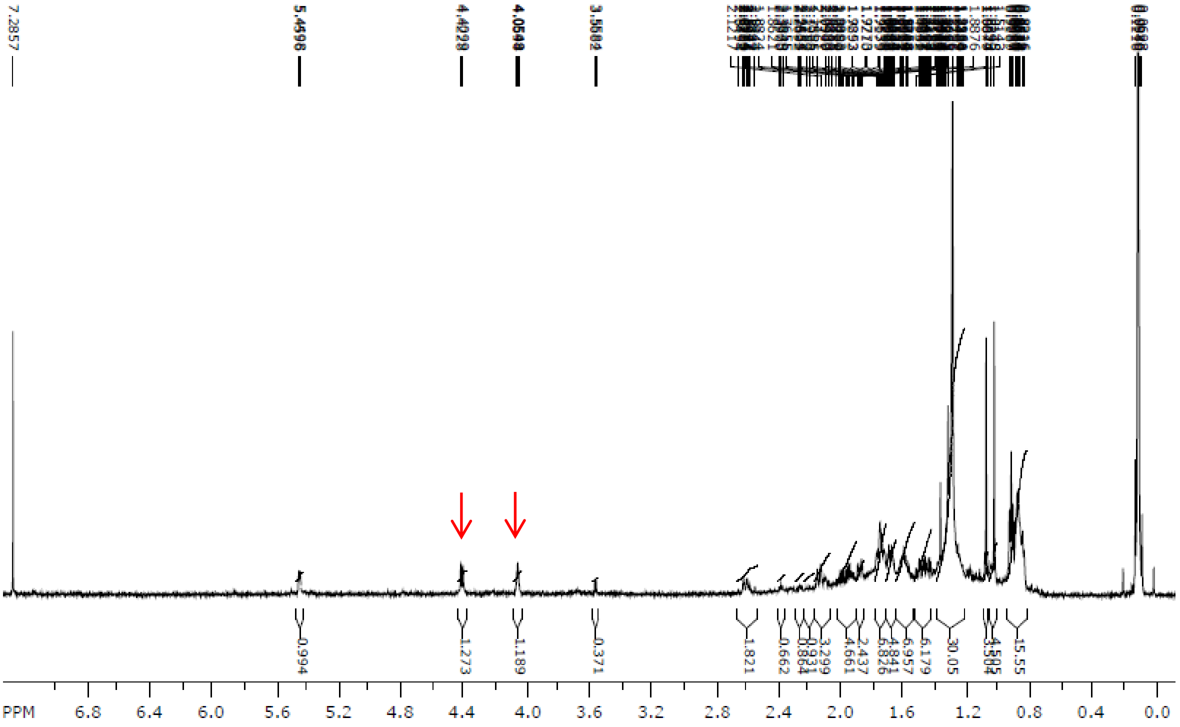

**B**

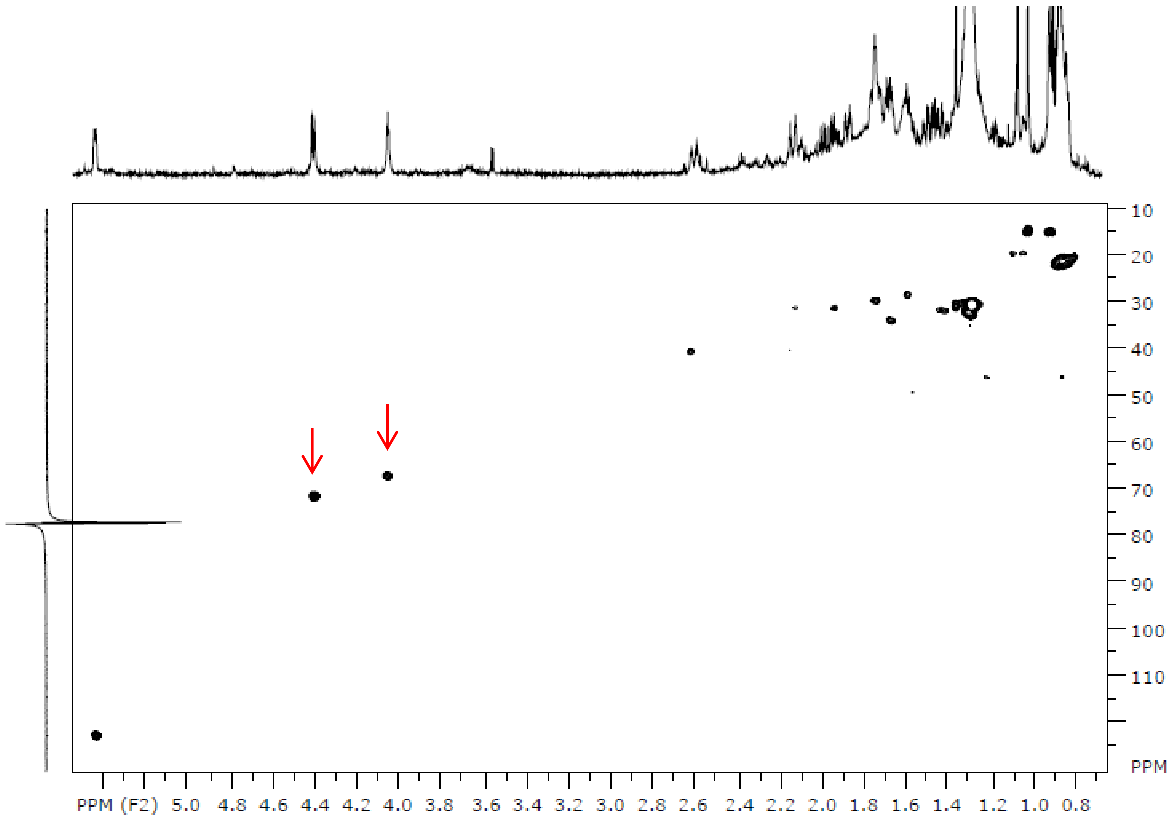


**Figure S17.** NMR spectra of the dihydroxy-3-deoxydehydroepiandrosterone product; **A**: $^{1}$H, **B**: HSQC.

#### d. Conversion of 5α-androstan-3-one (11) by CYP154C5 WT

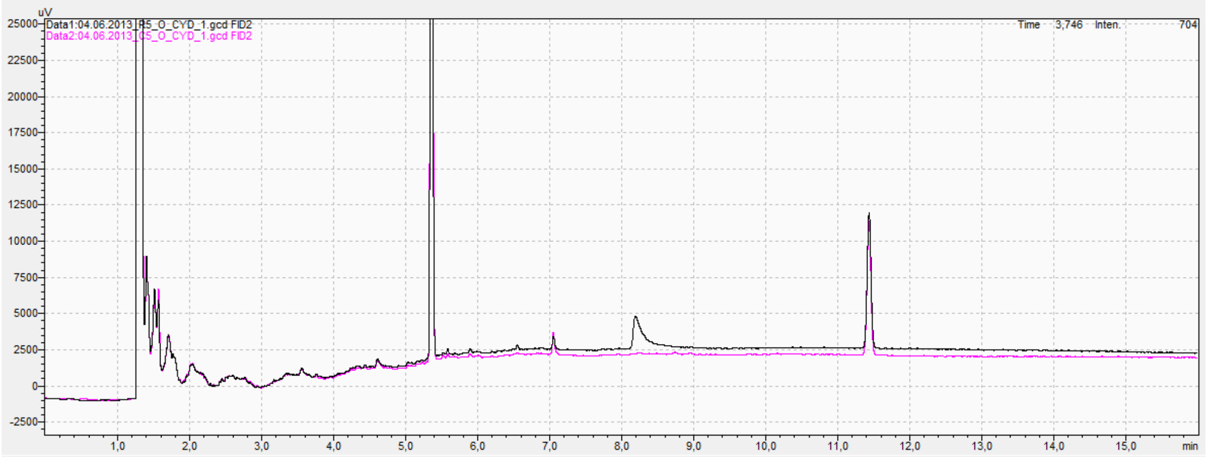


**Figure S18.** GC chromatogram (black) of 5α-androstan-3-one (**11**) conversion after 16 h catalysed by CYP154C5 wild type. The peak at 5.37 min corresponds to the substrate and the formed product eluted at 8.24 min. For comparison, the GC chromatogram of the respective control reaction (magenta) lacking CYP154C5 is shown as well.

**15α-hydroxy-5α-androstan-3-one** $^{1}$H-NMR (600 MHz, $CDCl_3$): δ 4.50 (1H, m, 15β-H), 2.45-2.23 (2H, m, 2α-H, 2β-H), 2.29 (1H, t, $J_t$= 14.3, 4α-H), 2.14-2.06 (2H, m, 4β-H, 16α-H), 2.04 (1H, dq, $J_q$= 6.8 $J_d$= 2.3, 1α-H), 1.77-1.53 (8H, m, 5-H, 7-H, 11-H, 12-H, 17α-H, 17β-H), 1.44-1.25 (7H, m, 1β-H, 6α-H, 6β-H, 8-H, 11-H, 12-H, 16β-H), 1.18 (1H, dd, $J_d$= 12.5 $J_d$= 5.9, 14α-H), 1.06-0.98 (4H, m, 7β-H, 19-CH3), 0.85 (1H, m, 9α-H), 0.75 (3H, s, 18-CH3). $^{13}$C-NMR (150 MHz, $CDCl_3$): δ 211.2 (3-C), 71.7 (15-C), 54.0 (9-C), 52 (14-C), 52 (16-C), 46.6 (5-C), 44.7 (4-C), 41.9 (13-C), 38.6 (1-C), 38.5 (2-C), 38.2 (12-C), 37.3 (17-C), 35.8 (10-C), 35.2 (8-C), 31.9 (7-C), 28.9 (6-C), 21.1 (11-C), 18.7 (19-C), 11.5 (18-C). Molecular weight: 290 g mol$^{-1}$; GC-MS: Retention time: 7.52 min. M$^{+}$: 290; m/z (%): 290 (11), 272 (41), 257 (65), 231 (76), 200 (50), 107 (57), 81 (71), 67 (74), 55 (100), 414 (93).

**A**

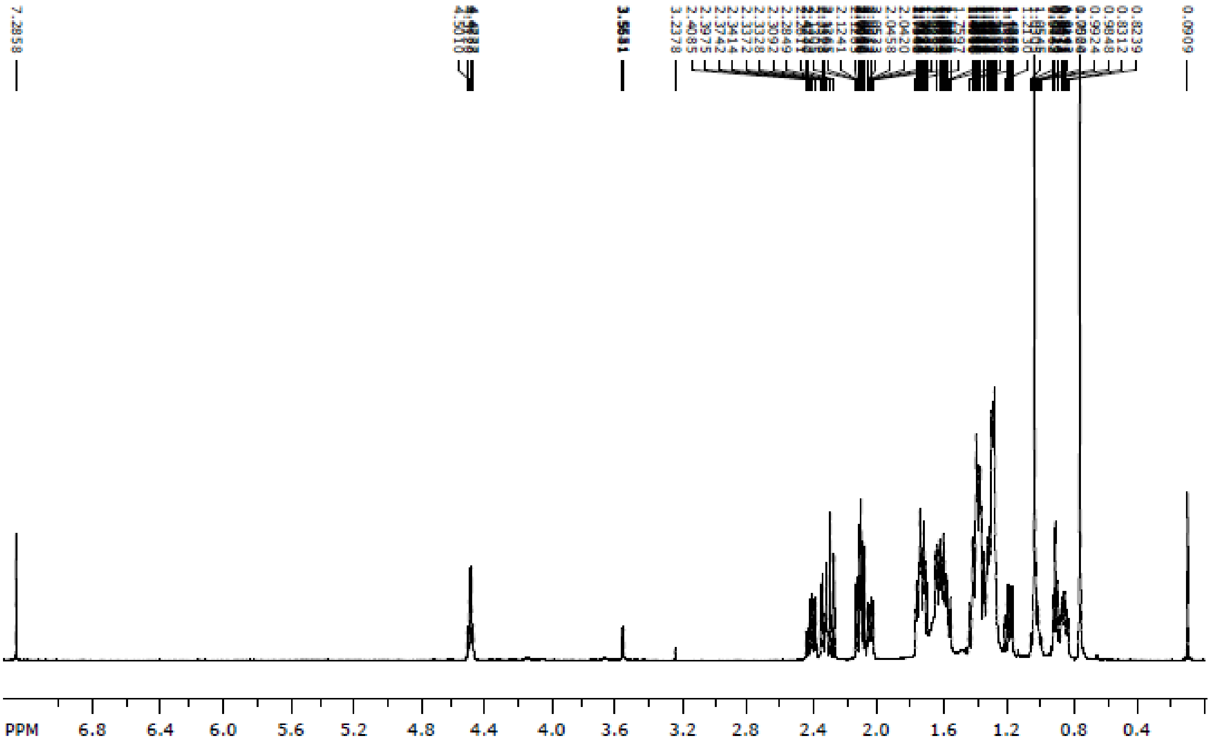

**B**

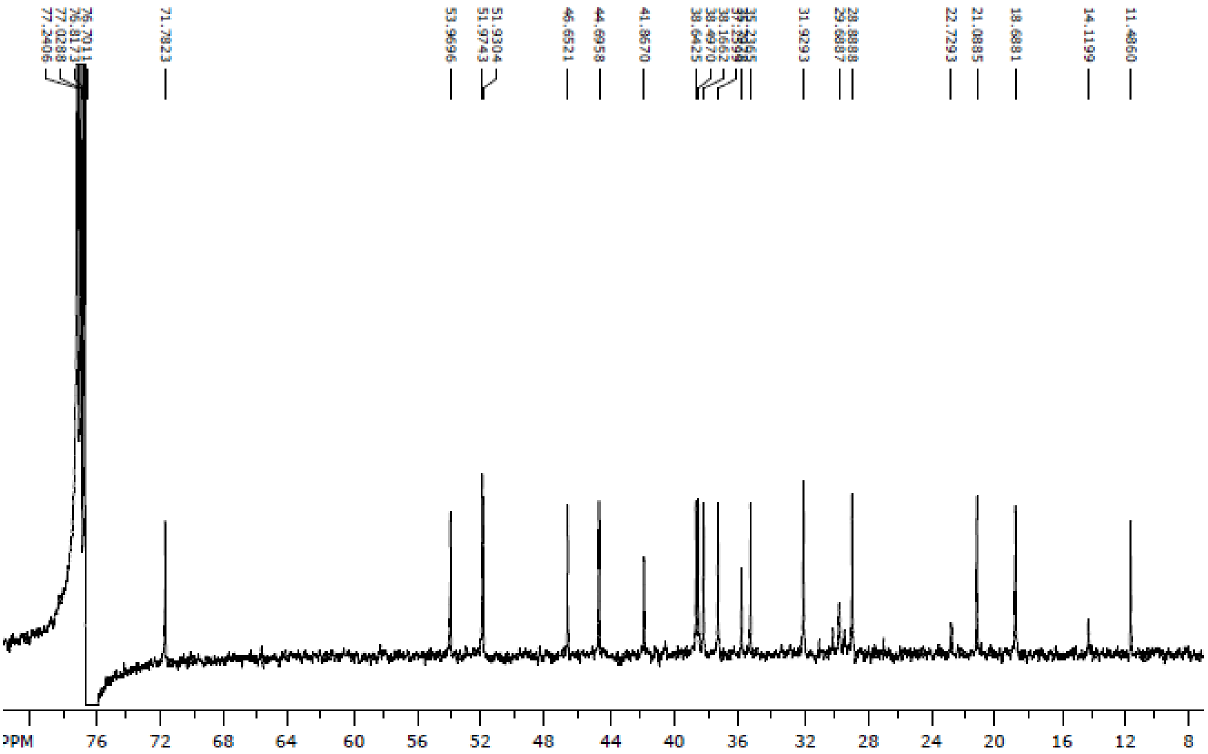

77.2406
77.0288
76.8173
76.7011
71.7823
53.9696
51.9743
51.9304
46.6521
44.6958
41.8670
38.6425
38.4970
38.1662
31.9293
29.6887
28.8888
22.7293
21.0885
18.6881
14.1199
11.4860
PPM 76 72 68 64 60 56 52 48 44 40 36 32 28 24 20 16 12 8


**C**

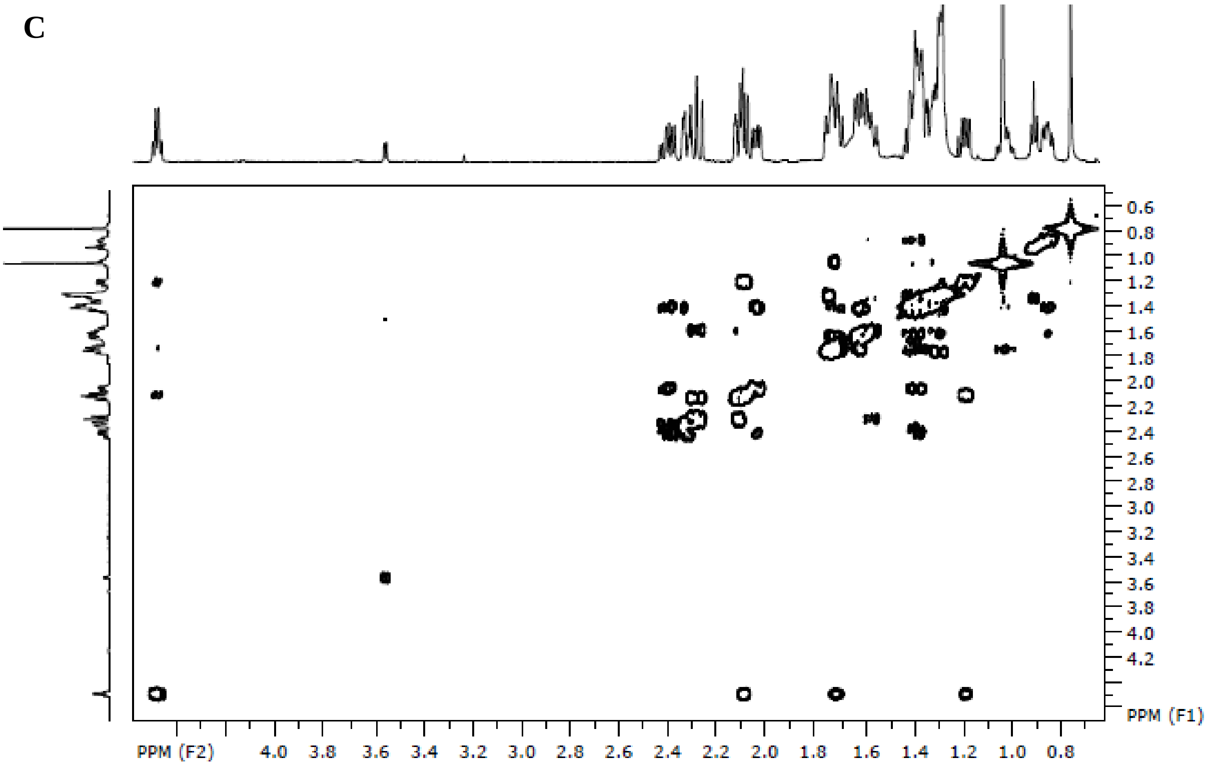

0.6 0.8 1.0 1.2 1.4 1.6 1.8 2.0 2.2 2.4 2.6 2.8 3.0 3.2 3.4 3.6 3.8 4.0 4.2
PPM (F1)
PPM (F2) 4.0 3.8 3.6 3.4 3.2 3.0 2.8 2.6 2.4 2.2 2.0 1.8 1.6 1.4 1.2 1.0 0.8

**D**

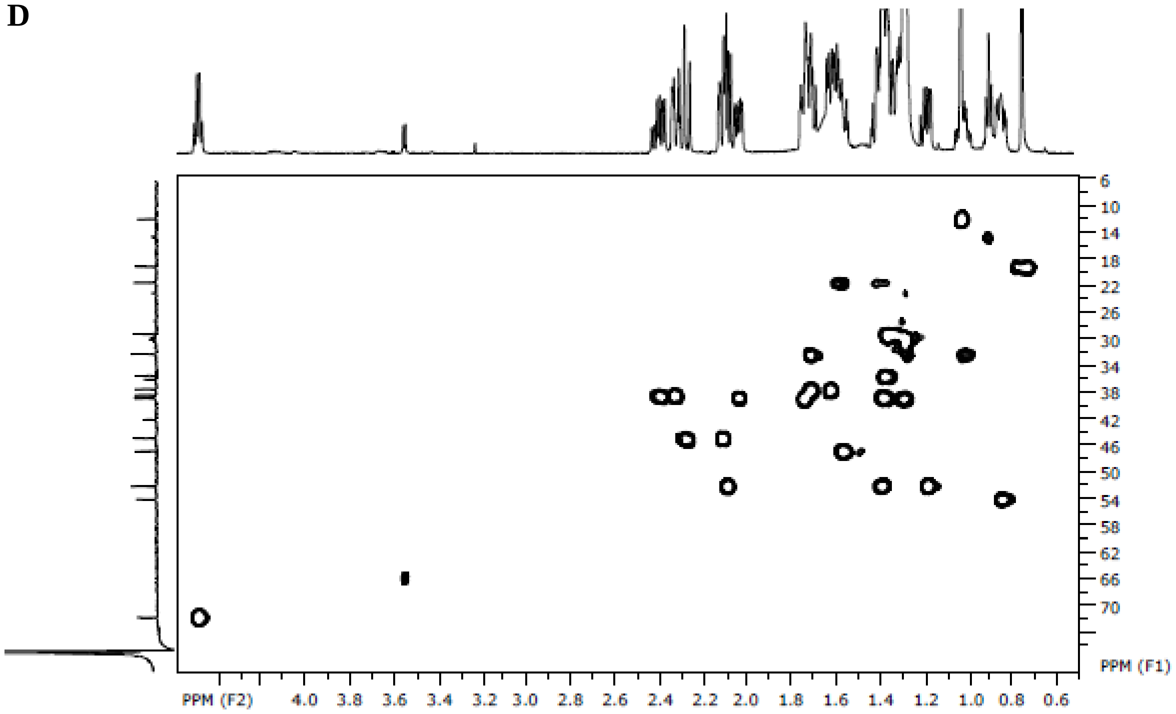


**E**

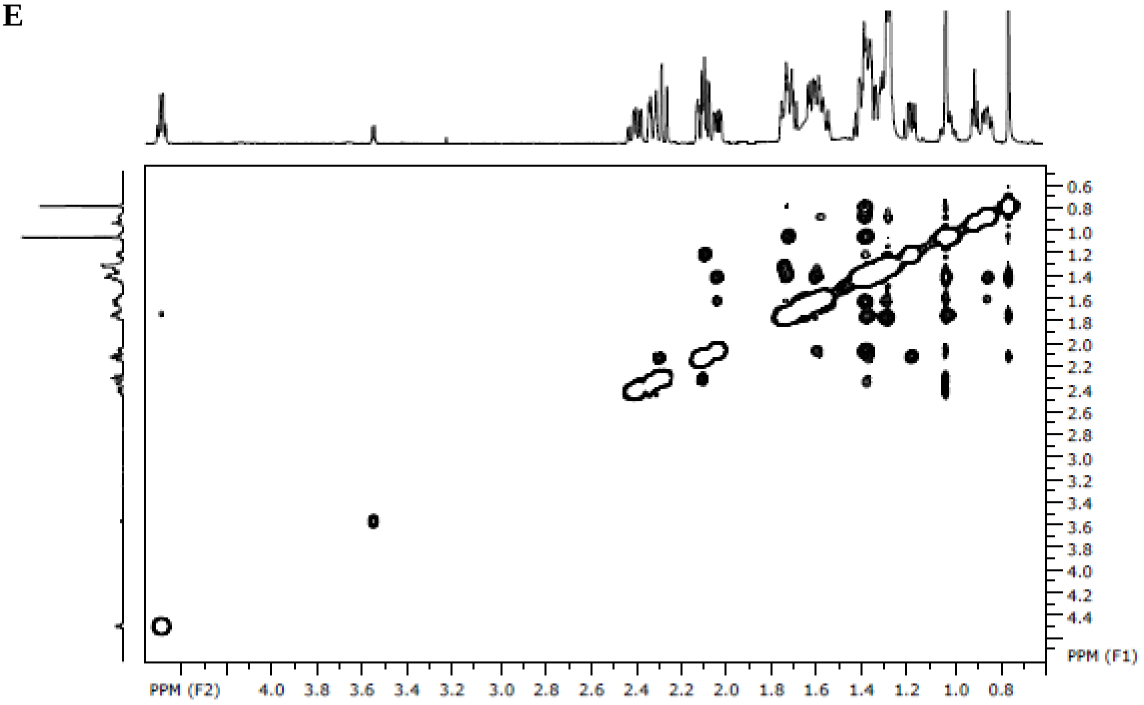


**Figure S19.** NMR spectra of 15α-hydroxy-5α-androstan-3-one; **A**: $^{1}H$, **B**: $^{13}C$, **C**: COSY, **D**: HSQC, **E**: NOESY.
